# Oscillating and stable genome topologies underlie hepatic physiological rhythms during the circadian cycle

**DOI:** 10.1101/2020.06.12.145771

**Authors:** Jérôme Mermet, Jake Yeung, Félix Naef

## Abstract

The circadian clock drives extensive temporal gene expression programs controlling daily changes in behavior and physiology. In mouse liver, transcription factors dynamics, chromatin modifications, and RNA Polymerase II (PolII) activity oscillate throughout the 24-hour (24h) day, regulating the rhythmic synthesis of thousands of transcripts. Also, 24h rhythms in gene promoter-enhancer chromatin looping accompany rhythmic mRNA synthesis. However, how chromatin organization impinges on temporal transcription and liver physiology remains unclear. Here, we applied time-resolved chromosome conformation capture (4C-seq) in livers of WT and arrhythmic *Bmal1* knockout mice. In WT, we observed 24h oscillations in promoter-enhancer contact at multiple loci including the core-clock genes *Period1, Period2* and *Bmal1*. In addition, we detected rhythmic PolII activity, chromatin modifications and transcription involving stable chromatin loops at gene promoters representing key liver function such as glucose and lipid metabolism and detoxification. Intriguingly, these contacts persisted in clock-impaired mice in which both PolII activity and chromatin marks no longer oscillated. Finally, we observed chromatin interaction hubs connecting neighbouring genes showing coherent transcription regulation across genotypes. Thus, both clock-controlled and clock-independent chromatin topology underlie rhythmic regulation of liver physiology.

## Introduction

Human behaviour and physiology have adapted to daily recurring inputs from the environment. Most animals including mammals have integrated a time monitoring device, known as the circadian clock, allowing them to resonate with these 24h external cues. The coupling of our internal clock with environmental day-night or light-dark cycle controls our wake-sleep rhythm but also, as illustrated here, the 3-dimensional (3D) shape of chromosomes in cells of the intact liver in mice. In mammals, the suprachiasmatic nucleus (SCN) receives light input from the retina and synchronizes peripheral organs through direct and indirect signalling (Dibner et al.). The circadian clock is molecularly encoded and relies on interlocked feedback loops of gene function ticking in virtually every cell of the body (Nagoshi et al.). In this model, BMAL1 and CLOCK transcription factors (TF) regulate the expression of their own repressors including *Period* (*Period1, Period2* and *Period3*) and *Cryptochrome* (*Cryptochrome1* and *Cryprochrome2*) genes (Takahashi). Clock-based and organ-specific TF activities interweave to regulate tissue-specific rhythms in transcriptional programs and physiology (Zhang et al., Yeung et al.). For example, in mouse liver, TF binding as well as chromatin modifications and accessibility and PolII activity fluctuate genome-wide and drive the rhythmic expression of thousand genes important for hepatic functions (Rey et al., Koike et al., Sobel et al.). Furthermore, rhythms in post-transcriptional mechanisms can drive oscillations in the abundance and activity of gene products (Wang et al.,Mermet et al., 2017,Mauvoisin et al.,Wong and O’Neill).

In this context, changes in chromatin topology along the 24h day emerge as a regulatory layer for temporal gene expression (Mermet et al., Yeung and Naef). In the mammalian cell nucleus, chromatin is organized in a hierarchical network of 3D structures (Lieberman-Aiden et al., Dixon et al.). Regulatory interactions between DNA sequences, for example through a promoter-enhancer looping mechanism, mostly occur in *cis* within ∼0.1 to few megabases (Mb) large topologically associating domains (TADs) (Dixon et al.,Sanyal et al.,Fulco et al., Vermunt et al.). In cultured cells, oscillatory chromatin contacts were reported only at large genomic scale, such as between a clock output gene and DNA sequences located on *trans* chromosomes (Aguilar-Arnal et al.) or with the nuclear lamina (Zhao et al.). However, the latter mechanism was not observed in the mouse liver (Brunet et al.). At a smaller genomic scale, promoter-enhancer loops in mouse tissues were shown to underlie temporal and tissue-specific gene transcription, for example through alternative promoter usage (Yeung et al.). In fact, in mouse liver, the conformation of chromatin was captured at two opposite time points of the day genome wide, reporting that changes in genomic interactions occurred mostly at the sub-TAD scale (Kim et al.). In addition, rhythms in promoter-enhancer looping were reported to resonate with transcriptional cycles in mouse tissues, with high contact frequency synchronized with active transcript synthesis (Mermet et al., Kim et al., Beytebiere et al.). Remarkably, oscillations in the formation of these loops were abolished in arrhythmic *Bmal1* KO animals, showing that the circadian clock sustained daily changes in genomic interactions (Mermet et al.). Furthermore, the deletion of the daily connected *Cryptochrome1* (*Cry1*) intronic enhancer element abolished the dynamics of the loop and perturbed the *Cry1* transcription cycle (by reducing the frequency of transcriptional bursts), and eventually led to a short period phenotype of mutant animals (Mermet et al.). These detailed analyses pointed out an important role of chromatin topology in the control of diurnal transcription. However, it is not known whether changes in chromatin architecture systematically accompany daily rhythms in transcription.

Here, we investigated temporal changes in chromatin conformation in livers of WT and *Bmal1* KO animals using 4C-seq. We identified 24h rhythms in promoter-enhancer looping synchronized with the expression of the core-clock genes *Bmal1, Period1* and *Period2*. Furthermore, we showed that promoters of clock output genes, representing key physiological properties of hepatocytes such as metabolite synthesis, detoxification and lipid and glucose metabolism, recruited surrounding elements resembling enhancers. Although PolII activity and chromatin marks oscillated at interacting DNA sites in WT livers, promoter-enhancer contact frequency was maintained at similar levels during day-time and night-time, corresponding to active and inactive transcription, respectively. This suggested that rhythmic transcription took place over a static and closed conformation of chromatin loops. Remarkably, in *Bmal1* KO animals, PolII activity and chromatin modifications no longer oscillated at these sites, while their interaction frequency remained stable over time and at a comparable level to WT, showing a clock-independent mechanism of DNA looping. Finally, we found a cluster of stable interactions linking a set of genes that were co-regulated across time and genotypes. Overall, these findings further our understanding on the role of chromatin architecture in circadian gene regulation in animals.

## Results

### Oscillating chromatin contacts accompany rhythmic gene transcription of core-clock repressors and activators

To elucidate the dynamics of chromatin architecture along the day-night cycle in mouse tissues, we performed 4C-seq experiments every 4h for 24h as in (Mermet et al.) (Methods). 4C-seq probes the interaction frequencies between one “bait” DNA fragment and the entire genome (Simonis et al.). We placed 4C-seq baits at gene promoters of central components of the mammalian molecular clock such as *Bmal1* (Takahashi). *Bmal1* gene is rhythmically expressed in WT mouse liver with mRNA abundance peaking around ZT22 (Fig. 1A) (ZT: Zeitgeber time; ZT0 corresponds to onset of lights-on; ZT12 corresponds to onset of lights-off). As reported for other genes (Mermet et al.), the 4C-seq contacts for *Bmal1* were highly enriched on the *cis* chromosome, especially within a 2Mb region surrounding the bait position. This region contained ∼50% of *cis* counts for all time points (Supplementary Table) and comprised most gene regulatory interactions (Sanyal et al.). 4C-seq data were then normalized and analyzed applying a locally weighted multilinear regression (LWMR) using a Gaussian window (sigma=2.5 kb) centered on each fragment for local smoothing (Mermet et al.) (Methods). Temporal analysis revealed that the *Bmal1* promoter rhythmically contacted a genomic region spanning from ∼40 kilobases (kb) to ∼75 kb downstream of the transcription start site (TSS), with the contact frequency peaking around ZT18-20 at multiple 4C-seq peaks (Fig. 1B,C). To characterize interacting regions, we integrated time-resolved chromatin immuno-precipitation followed by high-throughput sequencing (ChIP-seq) experiments targeting PolII and chromatin marks typical of enhancer regulatory elements such as H3K27ac and DNase1 hypersensitive sites (DHS) (Sobel et al.). As expected, PolII loading was rhythmic across the entire *Bmal1* gene body and peaked around ZT18-22 (Fig. 1D). Multiple 4C-seq interaction sites peaking at ZT20 coincided with chromatin regions marked by rhythmic regulatory activity. For example, a preferential ZT20 contact ∼45 kb downstream of the bait corresponded to a conserved intronic region marked by rhythmic H3K27ac histone acetylation and DNase1 hypersensitivity peaking around ZT20 (Fig. 1D). Another ZT20 4C-seq peak located near exon5 ∼73 kb downstream of the bait coincided with a ZT20 DNase1 signal (although weak) (Fig. 1C,D). These data suggested oscillating DNA loops between rhythmically active enhancers and the *Bmal1* promoter.

**Figure 1.**
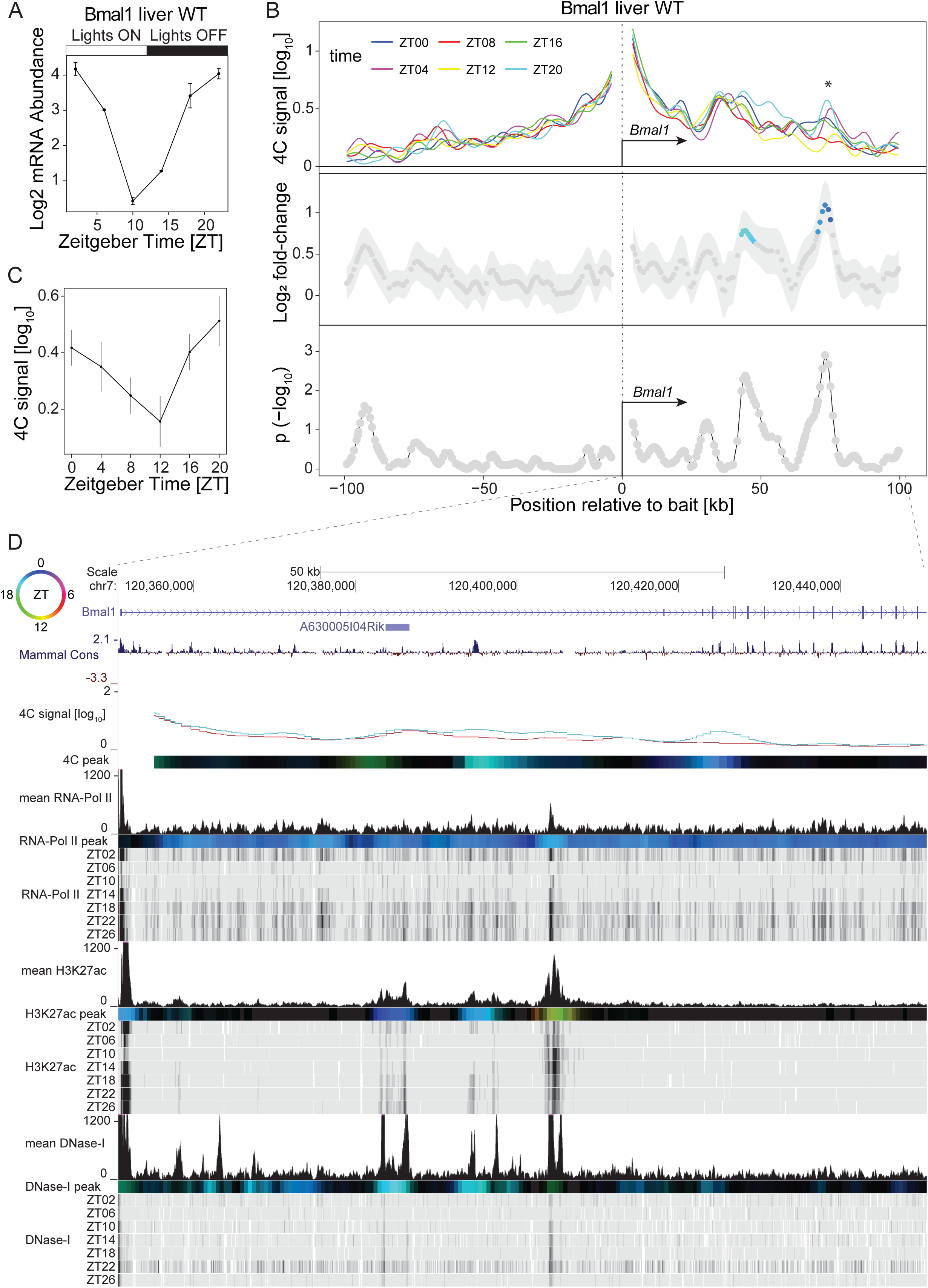
Time-resolved 4C-seq experiments revealed 24h rhythms in chromatin interactions at the *Bmal1* (*Arntl*) promoter in WT mouse liver. (A) *Bmal1* mRNA expression over time in WT mouse liver (Atger et al.). (B) 4C-seq signal over time from the *Bmal1* TSS bait in WT mouse liver (top panel) and log_2_ fold change (middle panel) and **−**log_10_(p) (lower panel, Methods) for rhythmicity analyses (Mermet et al.). Fragments with p<0.01 are colored according to peak time in contact frequency (color-coding as in following top left circle panel D). * = local maximum in differential genomic contact. n=1 in ZT04/ZT12/ZT20, n=2 in ZT00/ZT08/ZT16 (C) 4C-seq signal over time, adjacent to * (B). (D) 4C-seq signal at ZT20 (blue) and ZT08 (red), time-resolved ChIP-seq signal for PolII, H3K27ac and DNase1 hypersensitivity in WT mouse liver (Sobel et al.). Color-coded tracks represent peak time for 4C-seq signal and chromatin marks (see top left circle for time code, methods). Localized genomic regions marked by ZT18 to ZT00 H3K27ac and DNase1 hypersensitivity make chromatin contacts with the promoter region of *Bmal1* at ZT20.

Next, we explored chromatin architecture dynamics surrounding *Period1* (*Per1*) and *Period2* (*Per2*) genes that belong to the negative limb of the circadian molecular oscillator (Takahashi). *Per2* and *Per1* 4C-seq signals were largely enriched within a 2Mb window on *cis* chromosome (Supplementary Table 1). *Per2* mRNA is highest around ZT14 in WT liver (Fig. 2A). At the *Per2* locus, a large region extending from ∼35 kb to ∼70 kb downstream of the TSS contacted more frequently the promoter at ZT16, with two prominent oscillating contacts at 40 kb and 65 kb downstream of the TSS (Fig. 2B,C). The region 40 kb downstream of *Per2* TSS corresponded to multiple intragenic sites near the 3’ end of *Per2* containing rhythmic transcription and enhancer chromatin marks (PolII, H3K27ac, DHS) peaking around ZT16, as well as the *Hes6* gene in which PolII, H3K27ac and DHS peaked at ZT16 (Fig. 2D). The region 65 kb downstream of the TSS also contained H3K27ac and DHSs enhancer marks (Fig. 2D). Thus, these data showed 24h rhythms in enhancer-promoter contacts accompanying *Per2* transcription, and also dynamic gene-gene interactions with synchronised transcription. Furthermore, *Per1* mRNA is maximal around ZT10-ZT14 in WT mouse liver (Supplementary Fig. 1A). Two proximal regions contacted the *Per1* TSS more frequently at ZT12-ZT14 (Supplementary Fig. 1B). The first region was immediately upstream of the *Per1* TSS and corresponded to multiple sites with PolII, H3K27ac and DHSs (Supplementary Fig. 1C,D). The second site located ∼15-20 kb downstream of the TSS corresponded to the 3’ terminal region of *Per1* and coincided with H3K27ac and DHS sites, as well as the promoter region of *Hes7* (Supplementary Fig. 1C,D). These data suggest that these sites resembling enhancer elements diurnally contacted the promoter of *Per1*. We also noted a weak ZT02 preferential interaction 45 kb upstream the *Per1* TSS (Supplementary Fig. 1B).

**Figure 2.**
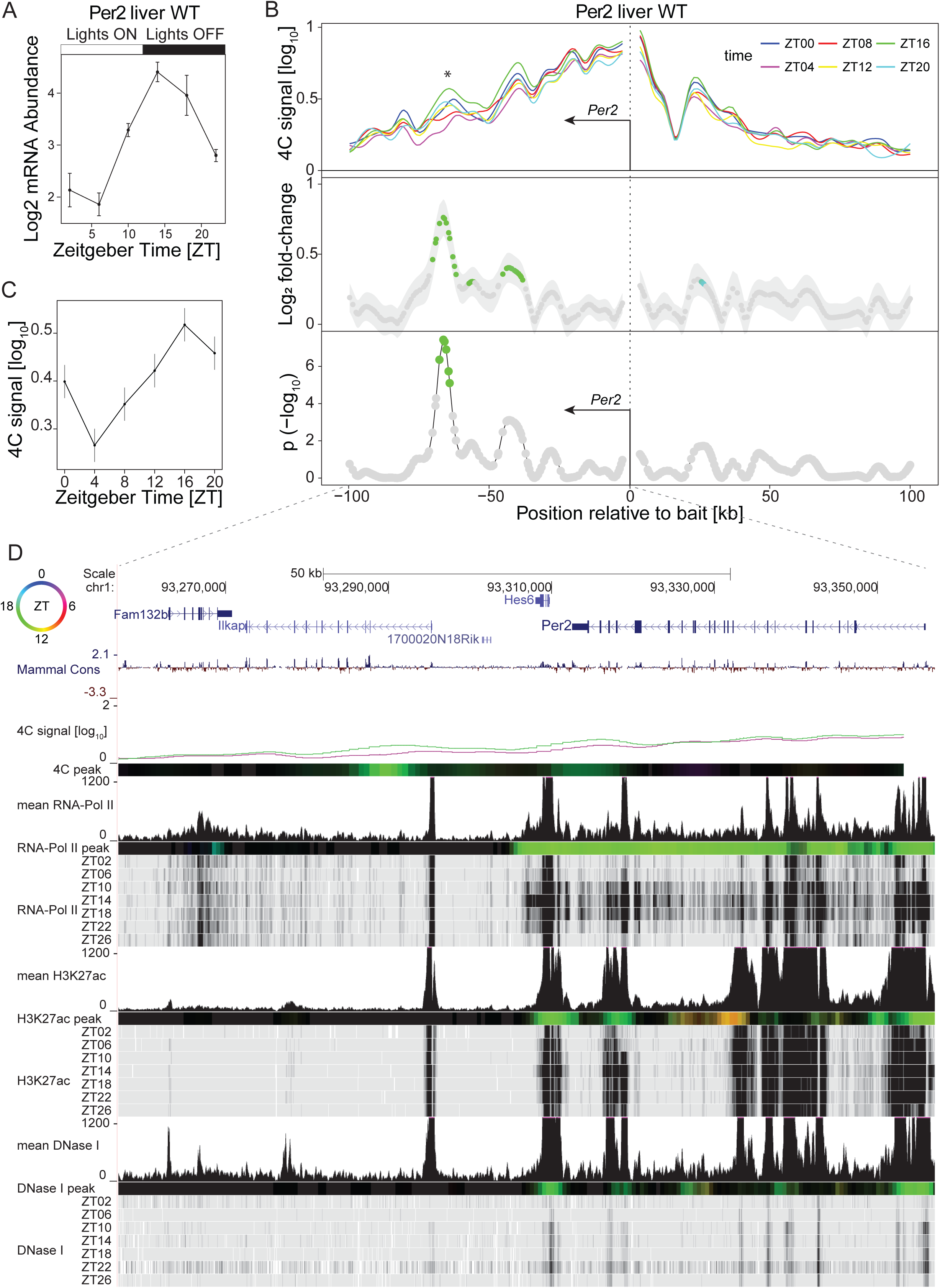
Time-resolved 4C-seq experiments revealed 24h rhythms in chromatin interactions of the *Period2* promoter in WT mouse liver. (A) *Period2* mRNA expression over time in WT mouse liver (Atger et al.). (B) 4C-seq signal over time from the *Period2* TSS bait in WT mouse liver (top panel) and log_2_ fold change (middle panel) and **−**log_10_(p) from rhythmicity analyses (Mermet et al.). Fragments with p< 0.01 are colored according to peak time in contact frequency (color-coding as in following top left circle panel D). * = local maximum in differential genomic contact. n=2. (C) 4C-seq signal over time adjacent to * (B). (D) 4C-seq signal at ZT04 (purple) and ZT16 (green), and time-resolved ChIP-seq signal for PolII, H3K27ac and DNase1 hypersensitivity in WT mouse liver (Sobel et al.). Colored tracks represent peak time for the 4C-seq signal and chromatin marks, (top left circle for peak-time color code, methods). The region interacting with the promoter of *Period2* at ZT16 coincided with multiple localized rhythmic signals in H3K27ac and DNase1 hypersensitivity peaking at ZT16.

Together, these time-resolved 4C-seq experiments revealed oscillating contacts between core-clock gene promoter and surrounding enhancers. The temporal dynamics of DNA interactions as well as chromatin features at enhancer sites were synchronized with the transcription of the target gene, with high contact frequency and enhancer activity coinciding with the peak time of mRNA synthesis.

### Stable DNA loops connect daily active enhancers with promoters of clock-ouput genes in mouse liver

Next, we explored chromatin interactions surrounding clock output genes. Two main transcriptional waves centered around ZT08 and ZT20 (Le Martelot et al.) underlie daily rhythms in the physiology of the mouse liver, including detoxification (Gachon et al.), glucose (Lamia et al.) and lipid metabolism (Aviram et al.), and metabolite synthesis (Nakahata et al., Ramsey et al., Peek et al.). Notably, clock-related TFs bind to the promoter of genes involved in carbohydrate and lipid metabolism (Rey et al., Zhang et al.). Thus, we selected promoters of rhythmically expressed genes involved in key physiological function of hepatocytes and profiled chromatin interactions at two time points (ZT08 and ZT20), which correspond to either the active or inactive transcription states of those genes in WT liver (Le Martelot et al.). *Nampt* encodes the nicotinamide phosphoribosyltransferase rate-limiting enzyme in the NAD biosynthesis pathway and BMAL1 binds to its promoter (Nakahata et al.,Rey et al.,Ramsey et al.). Moreover, the NAD+-dependent histone deacetylase SIRT1 inhibits CLOCK-BMAL1 TF activity (Ramsey et al.), illustrating the interlocking between the clockwork machinery and the metabolic state of the cell. *Nampt* transcripts accumulate rhythmically in the liver of WT mice and peak around ZT14 (Fig. 3A). Again, *Nampt* 4C-seq signals were mostly confined within the 2Mb region surrounding the bait at both time points (Supplementary Table 1). Unlike the dynamics observed for core clock genes, Z-scores were centered around zero across the 2Mb signal-rich region showing no differential interactions between ZT08 and ZT20 (Fig. 3B). In particular, major 4C-seq interaction sites stood out at 50 kb and 125 kb upstream of the bait position (Fig. 3B), suggesting that these regions contacted the promoter of *Nampt* at a similar frequency during day and night. Time-resolved ChIP-seq experiments showed rhythmic loading of PolII along the *Nampt* gene body peaking at ZT10, showing that the accumulation of the mature mRNA was delayed compared to the rhythm of transcript synthesis (Fig. 3A,C and D). The two upstream interacting regions were marked by rhythmic PolII loading (although the ChIP-seq signal is weak) and H3K27ac and DHS signals peaking at ZT10 (Fig. 3C,D). These data showed DNA loops connecting the promoter of *Nampt* with presumably daily active enhancers at similar a frequency between ZT08 and ZT20 in WT mouse liver. We also noted secondary 4C-seq peaks 200 kb upstream and 60 kb downstream of the *Nampt* bait at sites marked by DHS and weak H3K27ac.

**Figure 3.**
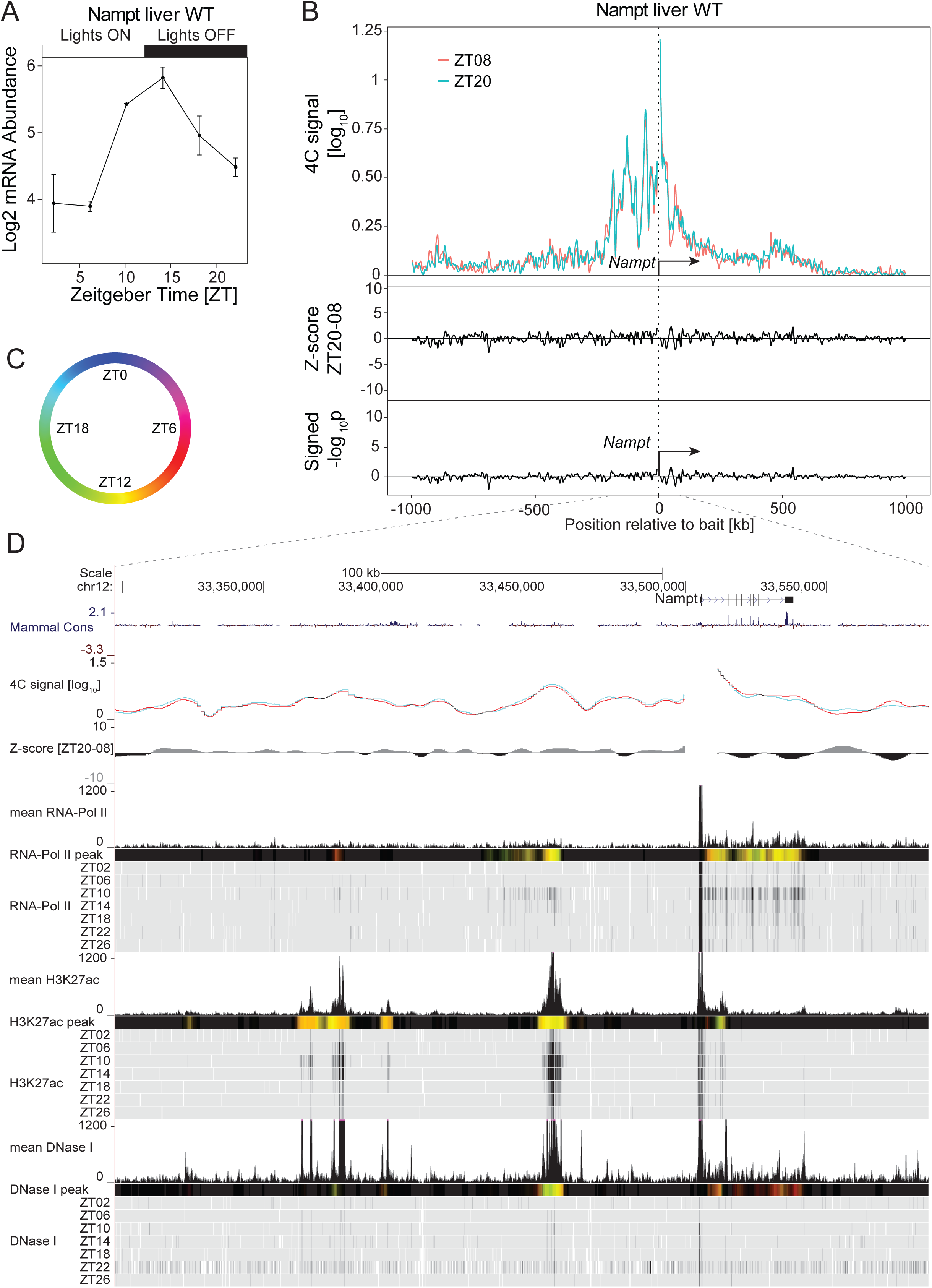
The *Nampt* promoter connects enhancer-like distal elements having rhythmic chromatin modifications. (A) *Nampt* mRNA accumulation over time in WT mouse liver from (Atger et al.). (B) 4C-seq signal at ZT08 (red, n=2) and ZT20 (green, n=4) in WT mouse liver in a 2Mb genomic window surrounding the *Nampt* bait (upper panel) and the corresponding Z-scores (middle track) and p-values (lower track). The most prominent 4C-seq peaks are located ∼50 kb and ∼125 kb upstream of the bait position. (C and D) Genome browser view with the *Nampt* 4C-seq signal at ZT08 (red) and ZT20 (blue) and time-resolved ChIP-seq signal for PolII, H3K27ac and DNase1 hypersensitivity in WT mouse liver (Sobel et al.) (D). Genomic regions located 50 kb and 125 kb upstream of the *Nampt* TSS are marked with DHSs and oscillating signals in H3K27ac peaking around ZT12, in sync with the transcription of *Nampt*. Colored tracks represent peak time in 4C-seq signal and chromatin mark (color code in C, methods).

A similar type of chromatin contact was observed at the locus of the clock-controlled gene *Lipg* (Yeung et al.). *Lipg* encodes a phospholipase protein involved in the degradation of lipoproteins and triglycerides (Riederer et al.). LIPG activity oscillates along the diurnal cycle in various tissues including the liver in rodents (Benavides et al.,Yeung et al.). *Lipg* transcripts accumulate rhythmically and peak at ZT14 in WT mouse liver (Fig. 4A). *Lipg* 4C-seq signals at ZT08 and ZT20 were enriched within the 2Mb genomic window in *cis*, and Z-scores across this region did not reveal differential interaction between the time-points (Fig. 4B). Several peaks of interaction were positioned near genomic regions located at 30 kb, 85 kb and 150 kb downstream of the bait position (Fig. 4B), suggesting that these sites were recruited to the *Lipg* promoter at a similar frequency during day and night. As expected, PolII activity was rhythmic across the *Lipg* gene body and peaked at ZT10, that is, 4h earlier than the mRNA peak (Fig. 4C,D). Again, this meant that ZT08 and ZT20 4C-seq time points captured chromatin conformation during active and inactive transcript synthesis, respectively. The site 30 kb downstream of the *Lipg* TSS corresponded to the 3’ terminal region of *Lipg* characterized by rhythmic PolII loading, H3K27ac and DNase1 sites peaking at ZT10 (Fig. 4C,D). Similarly, ZT10 peaks in PolII and H3K27ac marked the site 85 kb downstream of the TSS that contained multiple, although arrhythmic, DHSs. DHSs and traces of H3K27ac were observed at the interacting region 150 kb downstream of the TSS (Fig. 4C). Altogether, these data suggested chromatin contacts that were stable at the two time points corresponding to high and low transcription activity, and connecting the promoter of *Lipg* to downstream daily active enhancers.

**Figure 4.**
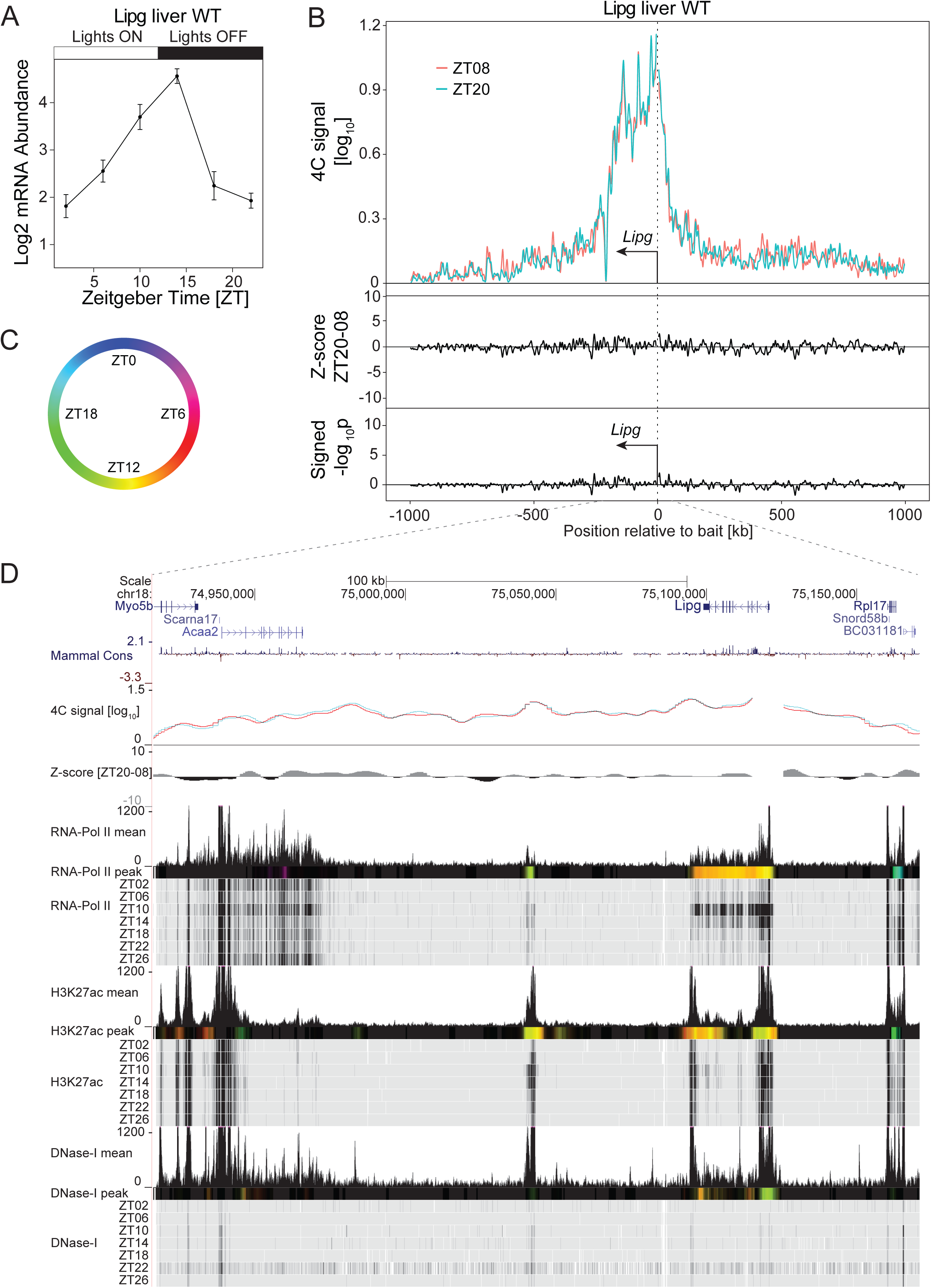
The *Lipg* promoter connects enhancer-like distal elements having rhythmic chromatin modifications. (A) *Lipg* mRNA accumulation over time in WT mouse liver (Atger et al.). (B) 4C-seq signal at ZT08 (red, n=3) and ZT20 (green, n=4) in WT mouse liver in a 2Mb genomic window surrounding the *Lipg* bait position (upper panel) and the corresponding Z-scores (middle track) and p-values (lower track). The most prominent 4C-seq peaks were located ∼30 kb, ∼85 kb and ∼150 kb downstream of the *Lipg* bait position. (C and D) Genome browser view with *Lipg* 4C-seq signals at ZT08 (red) and ZT20 (blue) and time-resolved ChIP-seq signal for PolII, H3K27ac and DNase1 hypersensitivity in WT mouse liver (D) (Sobel et al.). Genomic regions located 30 kb, 85 kb and 150 kb downstream of the *Lipg* TSS are marked by DHSs and rhythmic signals in H3K27ac peaking around ZT10-12, synchronously with the transcription of *Lipg*. Colored tracks represent peak time in 4C-seq signal and chromatin mark (color code in C, methods).

Consistently with the observations at the *Nampt* and *Lipg* loci, we found additional examples of rhythmically active enhancer-promoter pairs forming DNA loops that were stable during active and inactive transcription at genes essential for glucose (*Pfkfb3* and *Slc2a2*) and cholesterol metabolism (*Hmgcr*) in WT liver (Supplementary Figs. 2-5). This suggested that the clock-controlled transcriptional machinery can act over a frozen promoter-enhancer contact network that is resilient to transcriptional activity. Note that a scenario in which the dynamics in chromatin architecture would distinguish core-clock from clock output genes is, however, unlikely. Indeed, the promoter of the liver specific gene *Mreg* (Yeung et al.) clearly showed oscillations in the recruitment of intragenic regulatory sites in WT livers, with high contact frequency synchronized with active transcription (Supplemental Fig. 6). Thus, both stable and rhythmic promoter-enhancer loops underlie clock-output gene regulation in WT mouse liver.

### The *Nampt* and *Lipg* promoter-enhancer loops are maintained in *Bmal1* knock-out animals

Next, we asked if chromatin topology surrounding rhythmically transcribed gene promoters was maintained in clock-impaired animals. Indeed, we previously demonstrated that BMAL1-dependent chromatin topology controls the function of a distal regulatory enhancer targeting the *Cry1* gene in mouse tissues (Mermet et al.). To investigate if a functional clock also sustained static DNA loops connecting clock output gene promoters and enhancers, we measured chromatin conformations at the *Nampt* and *Lipg* loci in livers of *Bmal1* KO animals at ZT08 and ZT20. Both genes are expressed at lower and constant levels in livers of those arrhythmic animals compared to WT (Supplemental Fig. 7A and Supplemental Fig. 8A). Overall, the distributions of 4C-seq signals were comparable between time points and genotypes for both genes (Supplementary Table 1). Remarkably, the respective *Nampt* and *Lipg* promoter-enhancer pairs were maintained at similar levels in clock-impaired mice compared to WT and constantly from ZT08 to ZT20 (Supplemental Fig. 7B and 8B, respectively). Furthermore, for both genes, PolII loading and H3K27ac chromatin marks were overall arrhythmic and decreased at connected promoter-enhancer regions in *Bmal1* KO compare to WT, consistent with transcript profiles (Supplemental Fig. 7C,D and Supplemental Fig. 8C,D), with the exception of *Lipg* gene body where PolII rhythms were still detected. These data suggested that static promoter-enhancer DNA loops were maintained in a BMAL1-independent mechanism.

### A chromatin hub connects temporally co-transcribed genes

The above examples suggested that diurnal transcription could occur upon a static conformation of chromatin. As shown in other systems (Schoenfelder et al., Sexton et al.), in such a model, gene-gene interactions might allow transcriptional co-regulation. A remarkable example arguing in favor of this model was observed at the locus of the liver-specific and rhythmically expressed gene *Por* (Yeung et al.). *Por* encodes the cytochrome P450 oxidoreductase enzyme involved in the NADPH-dependent electron transport pathway; its rhythmic expression along the diurnal cycle contributes to detoxification in the mouse liver (Gachon et al.,Johnson et al.). In WT livers, ZT08 and ZT20 4C-seq experiments using the *Por* promoter as a bait revealed chromatin contacts with a region spanning from ∼50 kb upstream to ∼ 100 kb downstream of the bait position, showing multiple local peaks (Fig. 5B). The interaction frequency was similar between time points (Fig. 5B), as shown by the zero centered Z-scores in the *cis* region (Fig. 5B). The 50 kb region upstream corresponded to *Rhbdd2* gene, while the downstream interacting region coincided with the *Tmem120a, Mir7034l, Styxl1* and *Mdh2* genes (Fig. 5D). PolII loading peaked at ZT10 across the entire interacting locus, coinciding with a phase coherent accumulation of the transcripts around ZT10 (Fig. 5A,D), and validating that the ZT08 and ZT20 time points measured chromatin contacts during active and inactive transcription, respectively. H3K27ac signals marked all prominent interacting sites, showing rhythms at the *Por* and *Rhbdd2* genes, peaking around ZT10. These data suggest that genes with synchronous transcription cycles contacted each other in WT liver, and at constant frequency during day and night.

**Figure 5.**
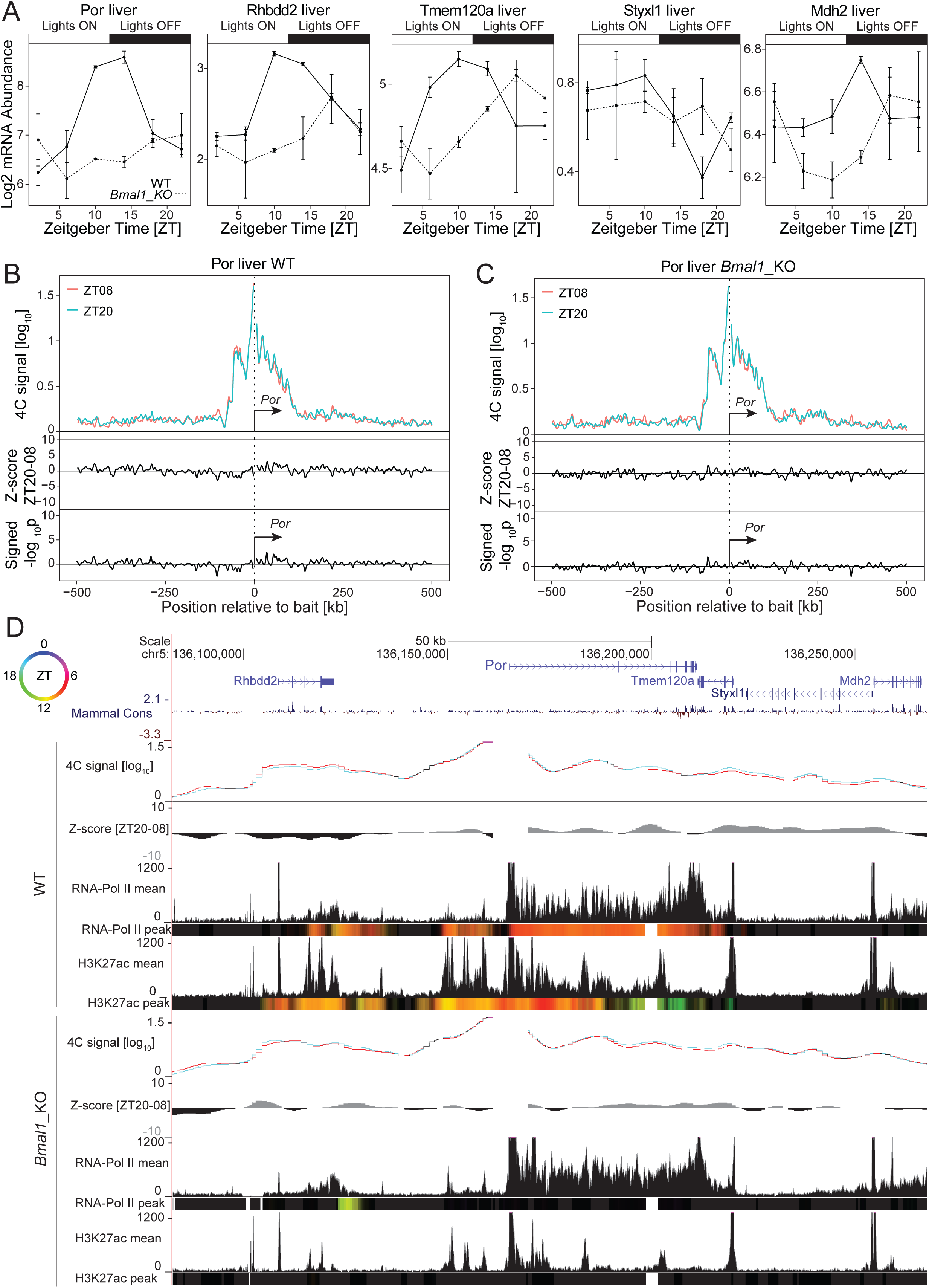
The *Por* gene stably connects surrounding genes having synchronized transcriptional dynamics in WT and *Bmal1* KO livers. (A) *Por, Rhbdd2, Tmem120a, Styxl1* and *Mdh2* transcripts accumulation over time in WT (solid line) and *Bmal1* KO (dashed line) livers (Atger et al.). (B and C) 4C-seq signal from *Por* TSS bait at ZT08 (red) and ZT20 (green) in a genomic window of 1Mb surrounding the bait in livers of WT (B, n=3 at ZT08 and n=4 at ZT20) and *Bmal1* KO (C, n=3) animals and the corresponding Z-scores and p-values. The most prominent interacting regions are located in the vicinity of *Por*. (D) Genome browser view of the *Por* 4C-seq signal at ZT08 (red line) and ZT20 (blue line) and the corresponding Z-scores in livers of WT and *Bmal1* KO animals. Mean and peak time of PolII and H3K27ac ChIP-seq signals are shown for both the WT and *Bmal1* KO conditions (Sobel et al.). Colored tracks represent peak time in chromatin mark (color code in the top left circle, methods). PolII loading and H3K27ac oscillate in WT livers at interacting regions, notably at *Por, Rhbdd2*, and *Tmem120a* genes and peak around ZT10, consistently with the rhythmic transcription of these genes. The dynamics of chromatin marks is lost over the entire locus in arrhythmic *Bmal1* KO livers, while 4C-seq signals are similar to WT conditions.

To further investigate the model of interaction between phase-coherent genes, we performed 4C-seq at the *Por* locus in livers of *Bmal1* KO mice. In these animals, the *Por* interacting sites were also connected at similar frequencies at ZT08 and ZT20, and at a comparable level to the WT conditions (Fig. 5C), suggesting that the chromatin architecture at this locus did not depend on a functional clock. Furthermore, *Por, Rhbdd2, Tmem120a, Styxl1* and *Mdh2* transcripts coherently accumulated at overall dampened, time delayed and lower levels in *Bmal1* KO compared to WT mice (Fig. 5A). Consistently, PolII loading and H3K27ac chromatin marks no longer oscillated across the entire interacting locus in livers of *Bmal1* KO compared to WT, and the level of histone acetylation was also reduced in clock-deficient animals (Fig. 5D). These data demonstrated that the *Por* interacting chromatin hub connected genes sharing similar temporal dynamics of transcription across the circadian cycle, and that the chromatin hub structure was resilient to transcription and clock-independent.

## Discussion

Here, we monitored chromatin contact dynamics across the 24h day at multiple core-clock and clock output gene promoters in livers of WT and arrhythmic *Bmal1* KO mice. By integrating temporal chromatin marks and transcriptome data, we aimed at characterizing the function of chromatin topology for temporal gene expression programs and physiology. Overall, we observed two classes of genomic interactions accompanying diurnal rhythms in gene expression.

In a first scenario, the promoter of rhythmically transcribed genes including the core-clock genes *Bmal1, Period1, Period2* (Fig. 1, Fig. 2 and Supplemental Fig. 1) and clock output genes such as *Mreg* (Supplemental Fig. 6 respectively) recruits surrounding elements in *cis* at a specific time of the day. These elements show rhythms in chromatin modifications typical of regulatory enhancers. Importantly, oscillations in both enhancer chromatin signatures and promoter-enhancer contact frequencies peaked in sync with the peak time of the target gene transcription (Supplementary figure 10). These findings agree with other works reporting promoter-enhancer looping coupled with rhythmic transcription activation (Mermet et al.,Kim et al.,Yeung et al.,Beytebiere et al.). We recently reported that rhythms in chromatin contact frequency depended on a functional clock (Mermet et al.), a mechanism that likely involves the recruitment of the mediator complex by clock TFs to connect distal sites (Mermet et al.,Yeung et al.,Kim et al.). Future work using arrhythmic animals might help at understanding, on a comprehensive scale, the role of the clock at modulating the 3-dimensional organization of DNA along the 24h day. Intriguingly, time-resolved 4C-seq assays also revealed that *Per2* contacted the immediately downstream TF *Hes6* at ZT16, and PolII activity within *Hes6* and *Per2* was also synchronized at ZT16 (Figure 2 and Supplementary figure 10). Furthermore, the paralogue *Per1* contacted the immediately downstream gene *Hes7* at ZT14, and *Hes7* mRNA accumulate synchronously in liver (Supplementary figure 1 and Supplementary figure 9). This observation suggests that duplicated genomic regions have conserved temporal dynamics of 3D chromatin architecture across the circadian cycle, and potentially allow transcriptional co-regulation of nearby genes. It would be exciting to challenge these hypotheses by comparing chromatin contact profiles of other duplicated genes within the clockwork machinery, such as *Cry1* and *Cry2* or *Bmal1* and *Bmal2*, among others.

In a second scenario, the promoter of rhythmically transcribed gene makes chromatin contacts with surrounding regulatory elements (e.g. *Nampt* and *Lipg*) or other genes (the *Por* interacting hub) in *cis*. Despite that contacting regions displayed rhythmic chromatin modifications (in the case of regulatory elements) and rhythmic transcription (in the case of gene-gene contacts), their relative contact frequency in the liver did not change across time, at least between the two time-points we probed (Fig. 3 and Fig. 4 and Supplemental Fig. 2-5). Since we captured chromatin conformation at two time points during the circadian cycle, changes in genomic interaction could occur at other time of the day, and would be missed in our experiments. However, the peak in transcription activity for most genes used as 4C-seq bait in this study, as reflected by PolII signal within gene bodies, reached maximum at ZT10 and was comparatively very low at ZT18-ZT22 (Fig. 3 and Fig. 4 and Supplemental Fig.2-5). Thus, the ZT08 and ZT20 4C-seq experiments captured genomic contacts of the genes analyzed here during active and inactive states of transcription, respectively. Furthermore, chromatin contacts persisted in clock-deficient animals, coinciding with a loss of rhythm in chromatin modifications at connected regions and in transcript synthesis (Fig. 5 and Supplemental Fig. 7,8). These data suggest a model in which gene promoter recruits distal regions, for example regulatory enhancers and/or co-regulated genes (Sexton et al.,Schoenfelder et al.), forming a chromatin hub structure, over which the clock machinery regulates rhythmic mRNAs synthesis (Xu et al.,Ghavi-Helm et al.) (Supplementary figure 10).

Although the data presented here (summarized in Supplementary figure 10) further our understanding of chromatin dynamics along day-night cycle, we acknowledge that 4C-seq revealed chromatin contacts for only a subset of gene promoters. To obtain a complete picture of the dynamics of chromatin architecture and its role in controlling circadian biology, it would be highly informative to perform promoter capture Hi-C (Javierre et al.) over time in WT and clock-impaired animals.

Finally, important finding from our previous work was that deleting an intronic enhancer rhythmically recruited to the *Cry1* gene promoter shortened the period of locomotor activity rhythm in animals (Mermet et al.). This effect propagated across regulatory layers, from the modulation of transcriptional bursting parameters to locomotor behavior. Here, we uncovered dozens of further non-coding regulatory elements potentially critical for the function of core-clock and clock output genes. It would be interesting to evaluate if multiple types of chromatin loops, for example stable versus dynamic, differentially affect transcriptional bursting parameters (Nicolas et al.,Bartman et al.). Furthermore, genetic manipulation might help appreciating more comprehensively the contribution of non-coding regulatory DNA to circadian biology, from transcription regulation to behavior.

## Material and Methods

### Animal housing

C57/BL6J (WT) and *Bmal1* KO animals were maintained at the EPFL animal house facility in 12hour/12hour light-dark cycle with 4 animals per cage. All experiments were approved by the Ethical Committee of the State of Vaud Veterinary Office (authorization VD3109).

### 4C-sequencing

4C-seq was performed in livers of 8 to 12 weeks old male with 4 biological replicates when comparing ZT08 versus ZT20 in WT animals and 3 biological replicates in *Bmal1* KO. 3 biological replicates were used in the 4h time-resolved 4C-seq experiments. Sample preparation and analyses were performed as in (Mermet et al.). In brief, livers were isolated and perfused with PBS before homogenization in 4 mL of 1×PBS including 1.5% formaldehyde for 10 minutes at room temperature. 25 mL of the following ice-cold buffer (2.2 M sucrose, 150 mM glycine, 10 mM HEPES at pH 7.6, 15 mM KCl, 2 mM EDTA, 0.15 mM spermine, 0.5 mM spermidine, 0.5 mM DTT, 0.5 mM PMSF) was added to the homogenates and kept for 5 min on ice. Homogenates were loaded on top of 10 mL of cushion buffer (2.05 M sucrose, 10% glycerol, 125 mM glycine, 10 mM HEPES at pH 7.6,15 mM KCl, 2 mM EDTA, 0.15 mM spermine, 0.5 mM spermidine, 0.5 mM DTT, 0.5 mM PMSF) and centrifuged at 10^5^ g for 45 minutes at 4°C. Nuclei were washed twice in 1× PBS and resuspended in 1 mL of 10 mM Tris-HCL (pH 8.0), 10 mM NaCl, 0.2% NP-40, and protease inhibitor cocktail (Complete Mini EDTA-free protease inhibitor cocktail; Sigma-Aldrich); kept for 15 minutes on ice; and washed twice with DpnII buffer (New England Biolabs). Thirty million nuclei were incubated for 10 minutes at 65°C in DpnII buffer and triton X-100 was added to 1% final concentration. Chromatin was digested overnight with 400 U of DpnII (New England Biolabs) at 37°C with shaking. Digested chromatin was then diluted in 8-mL of ligation buffer containing 3000 U of T4 DNA ligase for 4 h at 16°C plus 1 h at room temperature. 50 μL of 10 mg/mL proteinase K was added and samples were incubated overnight at 65°C. DNA was precipitated and resuspended in TE buffer (pH 8.0) containing RNase A, and incubated for 30 min at 37°C. Libraries were digested with NlaIII using 1U/μg of DNA template (New England Biolabs) overnight at 37°C. Digested products were ligated with 2000 U of T4 DNA ligase (New England Biolabs) for 4 h at 16°C in a 14-mL final volume. Circularized products were precipitated and resuspended in TE buffer (pH 8.0). Inverse PCRs were performed on 600 ng of circular DNA template per sample as described in (Mermet et al.). Inverse PCR primers are mentioned in the Supplemental Table 2.

### 4C-sequecing analysis

4C-seq data were analyzed as in (Mermet et al.). Briefly, demultiplexed read counts were mapped to the mouse genome (mm9) using HTSstation (David et al.). Samples were excluded from the analysis if more than 75% of restriction fragments did not have any count on a 2Mb region surrounding bait fragment (Supplementary table 1). The first five NlaIII fragments upstream and downstream of the bait were excluded in the analysis since they were suspected to be partially digested or self-ligated products. The 4C-seq signal was calculated using a locally weighted multilinear regression model (Mermet et al.). Fragment counts for each sample were normalized by the total fragments on the *cis*-chromosome (excluding the ten baits around the bait). To stabilize variance, the fragment counts *c* in each sample were log-transformed:

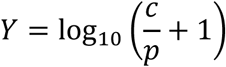

with p=500. For each position, the 4C-seq signals (*Y*) were modeled with fragment effects *a*_*i*_ and condition effects *b*_*j*_ (which can be time, tissue, or genotype). We estimated these effects by minimizing the weighted sum *S* of squared residuals across replicates *r:*

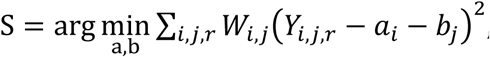, with weights *W*_*i,j*_ are defined as *W*_*i,j*_ = *w*_*g,i*_ × *w*_*s,j*_, where *w*_*g,i*_ is the Gaussian smoothing kernel (sigma=2500bp) at position *i*, and *w*_*S,j*_ a condition weight based on the number of samples with non-zero counts on fragment *i*. To compare between two conditions, we calculated p-values and condition effects at each genomic fragment using t-statistics. To detect rhythmic signal, we calculated the 24-hour Fourier coefficients of the condition effect (real and imaginary parts) from the six equally-spaced time points and used the chi-square test to test deviations from the null model that the real and imaginary parts have both a mean of zero.

### RNA-seq

Processed RNA-seq data were downloaded from (Atger et al.) (GSE73554) and rhythms were analyzed as in (Yeung et al.).

*H3K27ac and PolII ChIP-seq and DNase1-seq* data were downloaded from (Sobel et al.) (GSE60430) and analyzed as in (Mermet et al.). We binned the ChIP-seq and DNase1-seq signal (log2 counts per million) into 500 bp windows. We smoothed the signal by taking a running average across 7 bins (3 bins upstream and 3 bins downstream of the current bin). For each bin, we calculated the amplitude and phase of by fitting a harmonic regression model with a 24-hour period across the 7 bins. The rhythmic signal (amplitude and phase) was mapped to a color using hue (time of maximum signal), saturation (set to 1), and value (increased with increasing statistical significance) color scheme.

To obtain smooth color transitions, *v* was calculated using a Hill function with Hill coefficient *n* = 5 and 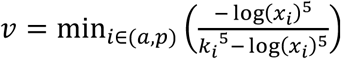, with *k*_*a*_ = 0.5, *k*_*p*_ = 4.5 and *x*_*a*_, *x*_*p*_ being amplitude and –log_10_(p) of the harmonic regression fit.

## Acknowledgements

We thank Frédéric Gachon for helpful discussions. This work was supported by the Swiss National Science Foundation Grants 31003A-153340 and 310030_173079 (to F.N.), a European Research Council Grant ERC-2010-StG-260667 (to F.N.), and the EPFL.

## Author contributions

J.M., J.Y. and F.N. conceived the study. J.M. performed the investigations. J.Y. and F.N. performed the analyses. J.M., J.Y. and F.N. wrote and reviewed the manuscript.

## Declaration: Data and code availability

The data is available on Gene Expression Omnibus (GSE139195). Analysis code and processed data can be explored through the Circadian4Cseq R package, which is available on github (https://github.com/jakeyeung/Circadian4Cseq). Reviewers can access the data using the secure token yfurumgedtwrnux.

## Supplemental Figure legends

**Supplemental Figure 1.**
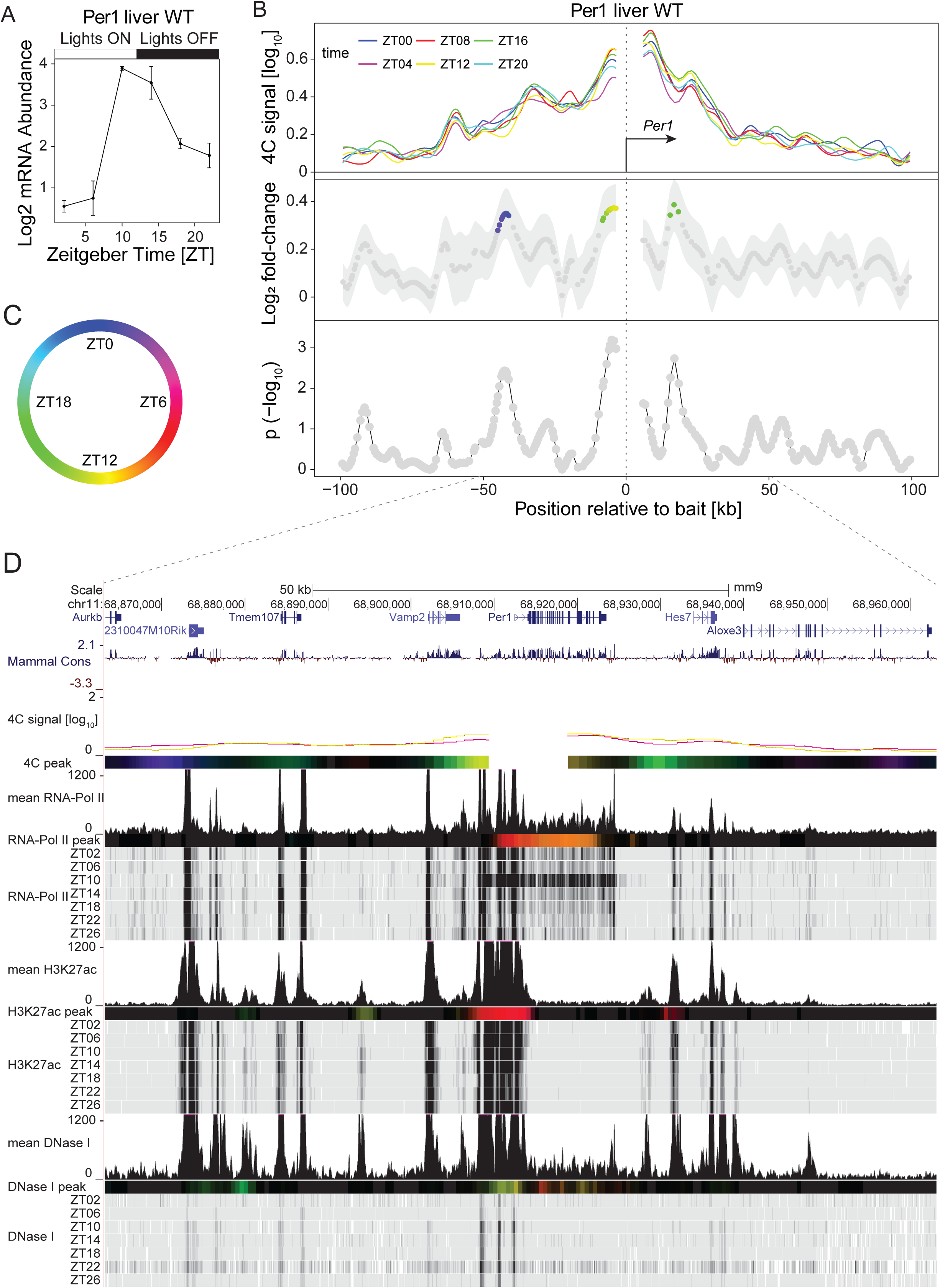
Time-resolved 4C-seq using the *Period1* TSS as bait revealed 24h rhythmic chromatin interactions in WT mouse liver. (A) *Period1* mRNA expression over time in WT mouse liver (Atger et al.). (B) 4C-seq signal over time for the *Period1* TSS bait in WT mouse liver (top panel) and log_2_ fold-change (middle panel) and **−**log_10_(p) (lower panel, methods) from rhythmicity analyses (Mermet et al.) (n=2). Although the 4C-seq signals were weak for the *Period1* bait, two localized genomic regions at ∼6 kb upstream and 17 kb downstream of the bait position were recruited preferentially at ZT12 to the *Period1* promoter. Fragments with p< 0.01 are colored by peak time in contact frequency as in following color coding (C). (D) *Period1* 4C-seq signal at ZT04 (purple) and ZT12 (yellow) and time-resolved ChIP-seq signals for PolII, H3K27ac and DNase1 hypersensitivity in WT mouse liver (Sobel et al.). Colored tracks represent peak time in 4C-seq signal and chromatin marks following the color code as in (C) (methods). The regions located immediately upstream and 17 kb downstream of the bait coincided with multiple localized peaks in H3K27ac and DNase1 hypersensitivity.

**Supplemental Figure 2.**
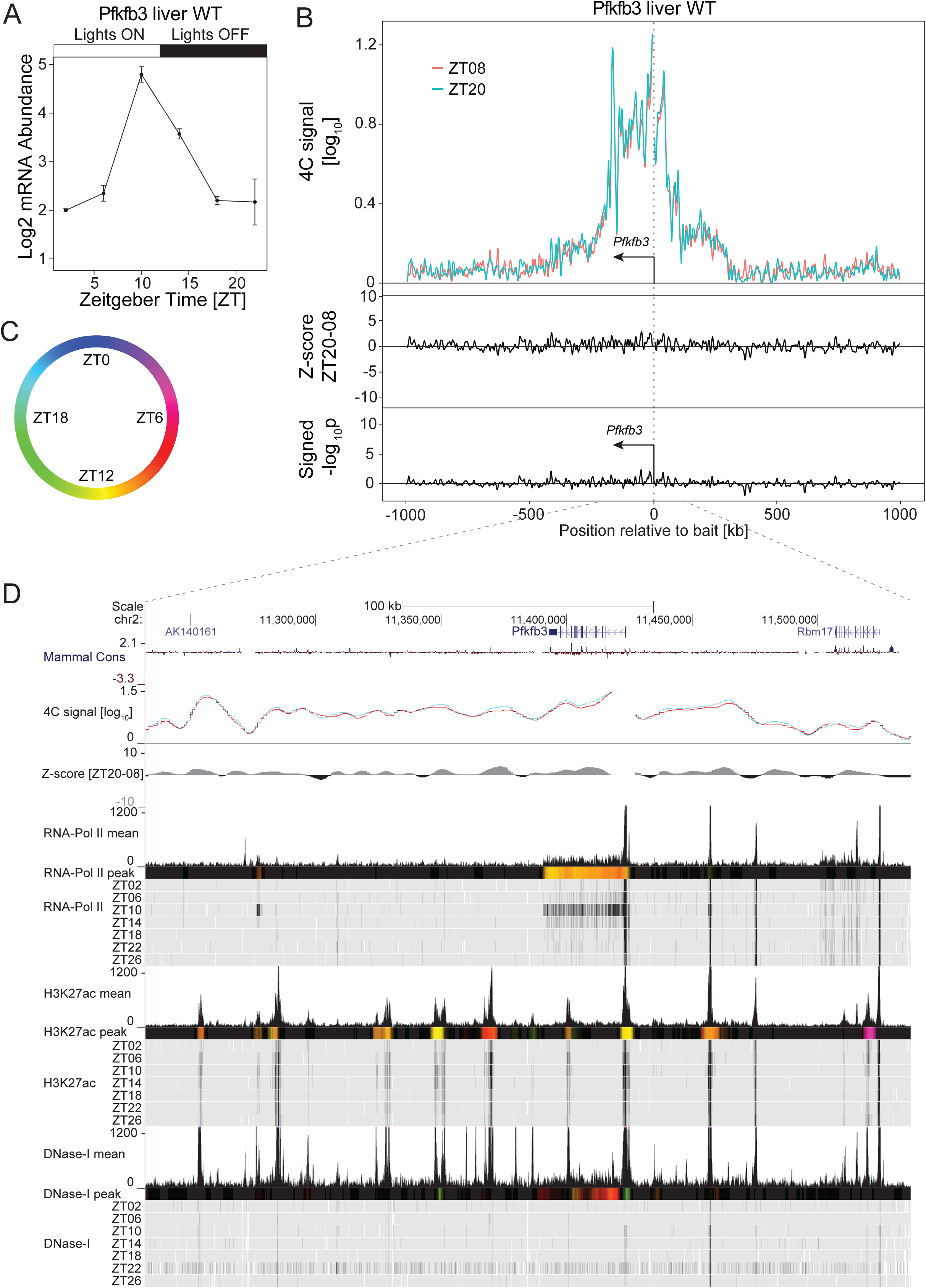
The promoter of *Pfkfb3* connects enhancer-like distal elements showing rhythmic chromatin modifications. (A) *Pfkfb3* mRNA expression over time in WT mouse liver (Atger et al.). (B) 4C-seq signal at ZT08 (red, n=3) and ZT20 (green, n=4) in WT mouse liver in a 2Mb genomic window surrounding the *Pfkfb3* bait position (upper panel) and the corresponding Z-scores (middle track) and p-values (lower track) revealing multiple prominent 4C peaks. (C and D) Genome browser viewing with *Pfkfb3* 4C-seq signal at ZT08 (red) and ZT20 (blue) and time-resolved ChIP-seq signal for PolII, H3K27ac and DNase1 hypersensitivity in WT mouse liver (D) (Sobel et al.). Colored tracks represent peak time in chromatin marks following the color code as in (C) (methods). Multiple regions interacting with the *Pfkfb3* TSS are marked by DHSs and rhythmic H3K27ac signals peaking around ZT10, synchronously with *Pfkfb3* transcription.

**Supplemental Figure 3.**
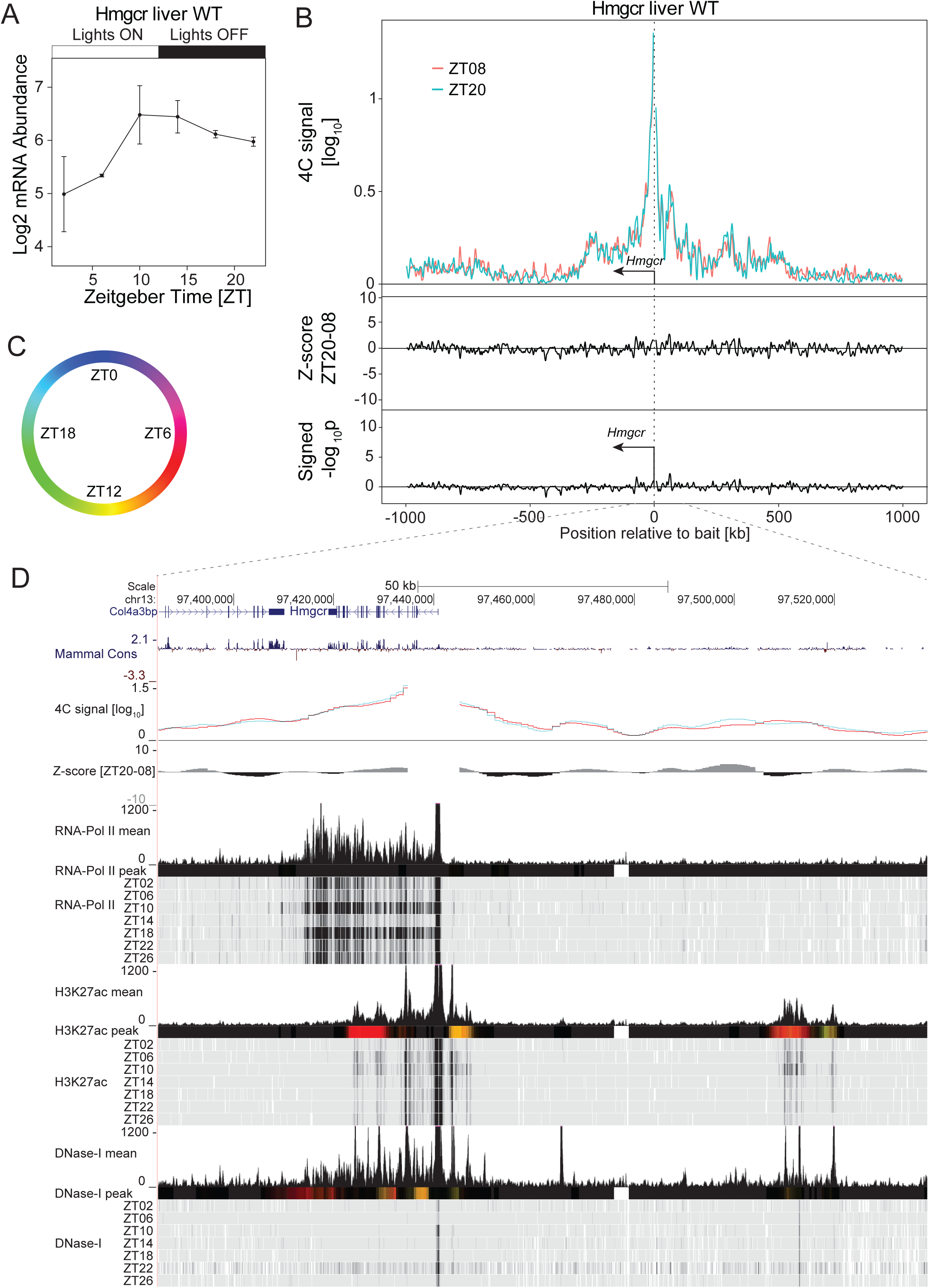
The promoter of *Hmgcr* connects enhancer-like distal elements showing rhythmic chromatin modifications. (A) *Hmgcr* mRNA expression over time in WT mouse liver (Atger et al.). (B) 4C-seq signal at ZT08 (red, n=2) and ZT20 (green, n=4) in WT mouse liver in a 2Mb genomic window surrounding the *Hmgcr* bait position (upper panel) and the corresponding Z-scores (middle track) and p-values (lower track) revealing local prominent 4C-seq peaks (although with low 4C-seq signals) located ∼70 kb upstream of the bait position. (C and D) Genome browser view with *Hmgcr* 4C-seq signal at ZT08 (red) and ZT20 (blue) and time-resolved ChIP-seq signal for PolII, H3K27ac and DNase1 hypersensitivity in WT mouse liver (D) (Sobel et al.). The genomic region 70 kb upstream of the *Hmgcr* TSS is marked by DHSs and rhythmic H3K27ac signals peaking around ZT08, coinciding with *Hmgcr* transcription time. Colored tracks represent peak time in chromatin marks following the color code as in (C) (methods).

**Supplemental Figure 4.**
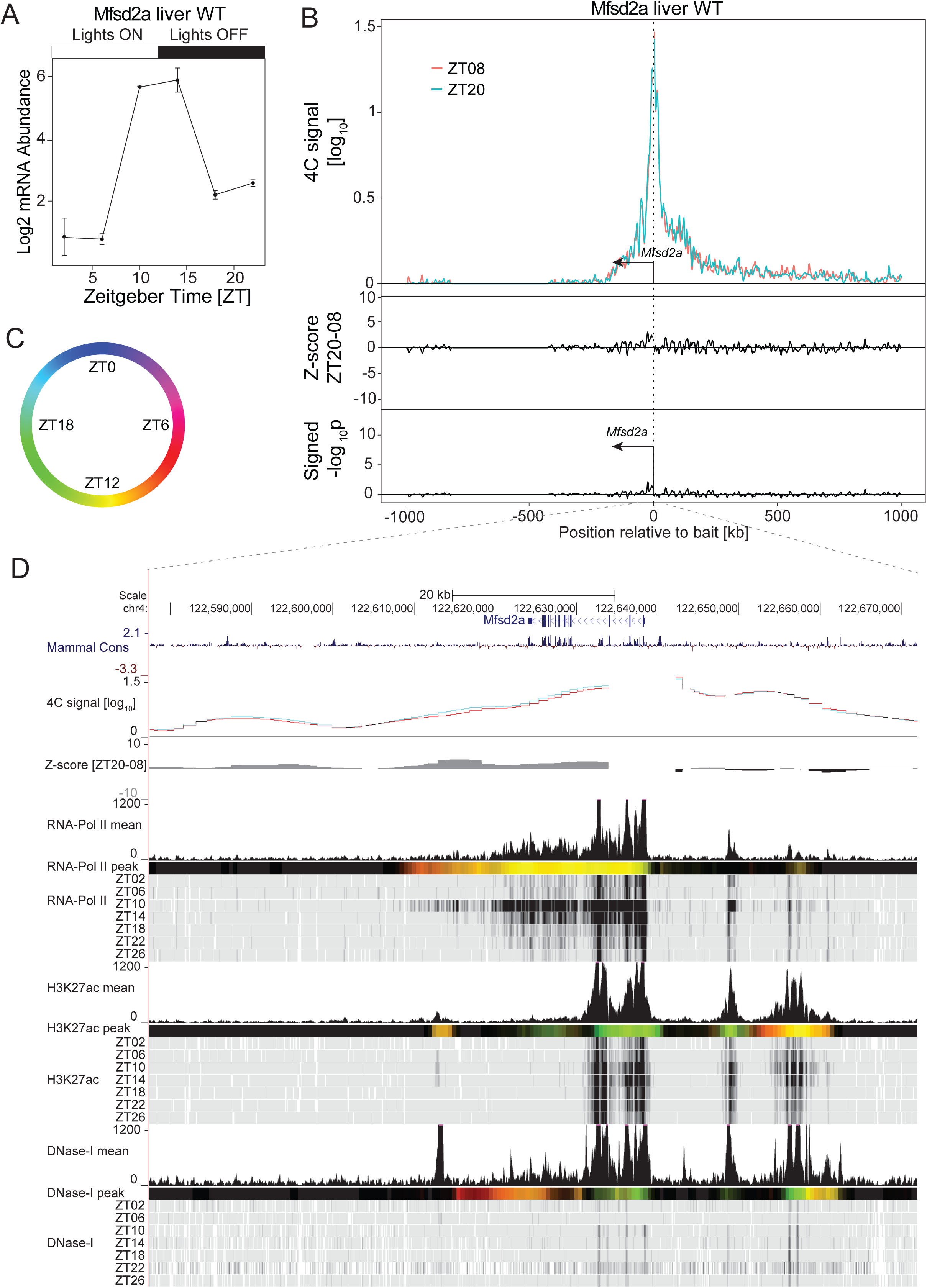
The promoter of *Mfsd2a* connects enhancer-like distal elements showing rhythmic chromatin modifications. (A) *Mfsd2a* mRNA expression over time in WT mouse liver (Atger et al.). (B) 4C-seq signal at ZT08 (red, n=3) and ZT20 (green, n=3) in WT mouse liver in a 2Mb genomic window surrounding the *Mfsd2a* bait position (upper panel) and the corresponding Z-scores (middle track) and p-values (lower track) revealing a localized prominent 4C-seq peak (although with low 4C-seq signals) ∼15 kb upstream of the bait position. (C and D) Genome browser viewing with *Mfsd2a* 4C-seq signal at ZT08 (red) and ZT20 (blue) and time-resolved ChIP-seq signal for PolII, H3K27ac and DNase1 hypersensitivity in WT mouse liver (D) (Sobel et al.). The genomic region located ∼15 kb upstream of the bait position is marked by DHSs and rhythmic H3K27ac signals peaking around ZT12, consistently with *Mfsd2a* transcription. Colored tracks represent peak time in chromatin marks following the color code as in (C) (methods).

**Supplemental Figure 5.**
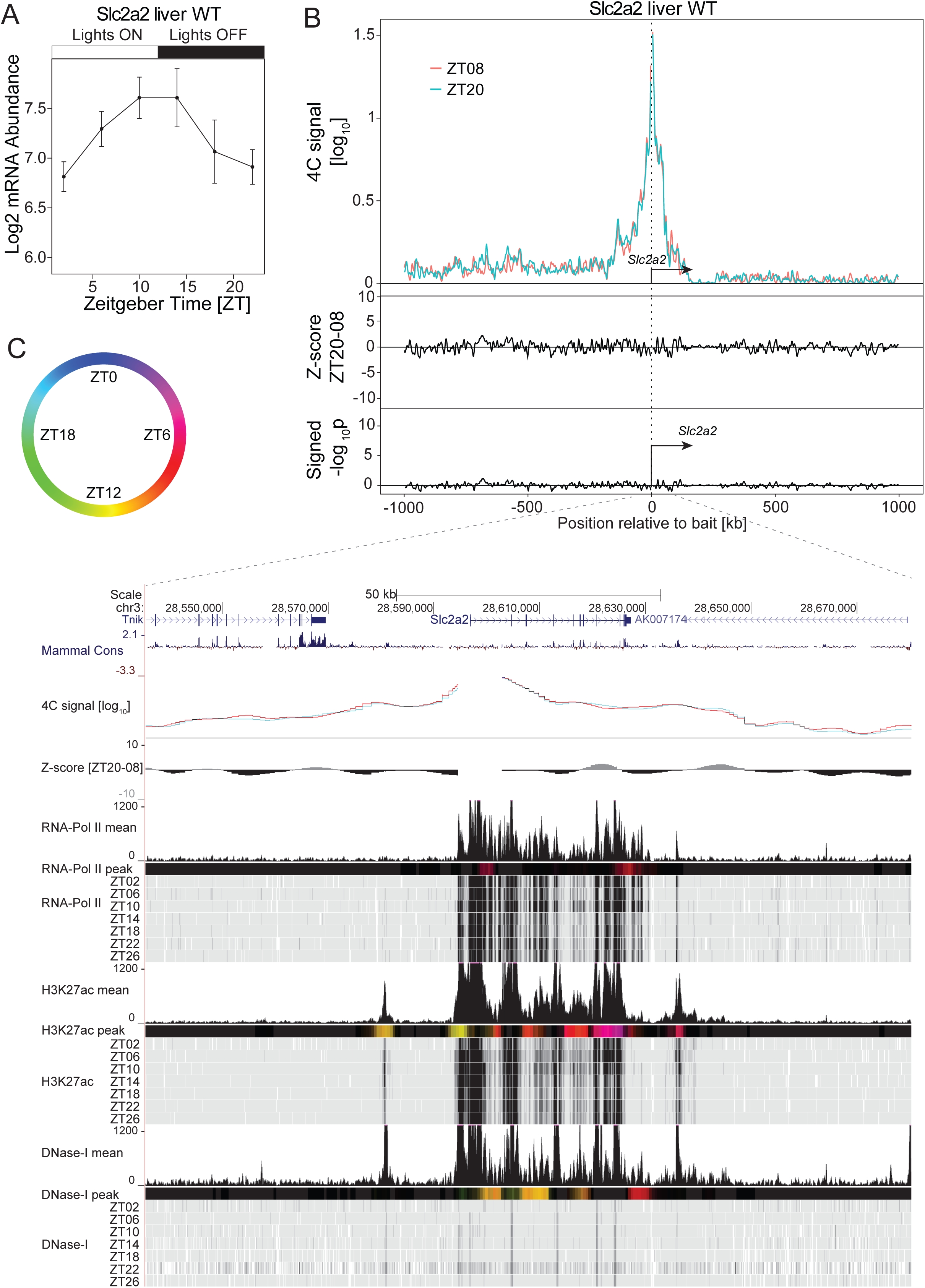
The promoter of *Slc2a2* connects enhancer-like distal elements. (A) *Slc2a2* mRNA expression over time in WT mouse liver (Atger et al.). (B) 4C-seq signal at ZT08 (red, n=2) and ZT20 (green, n=4) in WT mouse liver in a 2Mb genomic window surrounding the *Slc2a2* bait position (upper panel) and the corresponding Z-scores (middle track) and p-values (lower track) revealing multiple prominent 4C-seq peaks. (C and D) Genome browser viewing with the *Slc2a2* 4C-seq signal at ZT08 (red) and ZT20 (blue) and time-resolved ChIP-seq signal for PolII, H3K27ac and DNase1 hypersensitivity in WT mouse liver (D) (Sobel et al.). Regions interacting with the TSS of *Slc2a2* are marked by DHSs and H3K27ac peaks. Colored tracks represent peak time in chromatin marks following the color code as in (C) (methods).

**Supplemental Figure 6.**
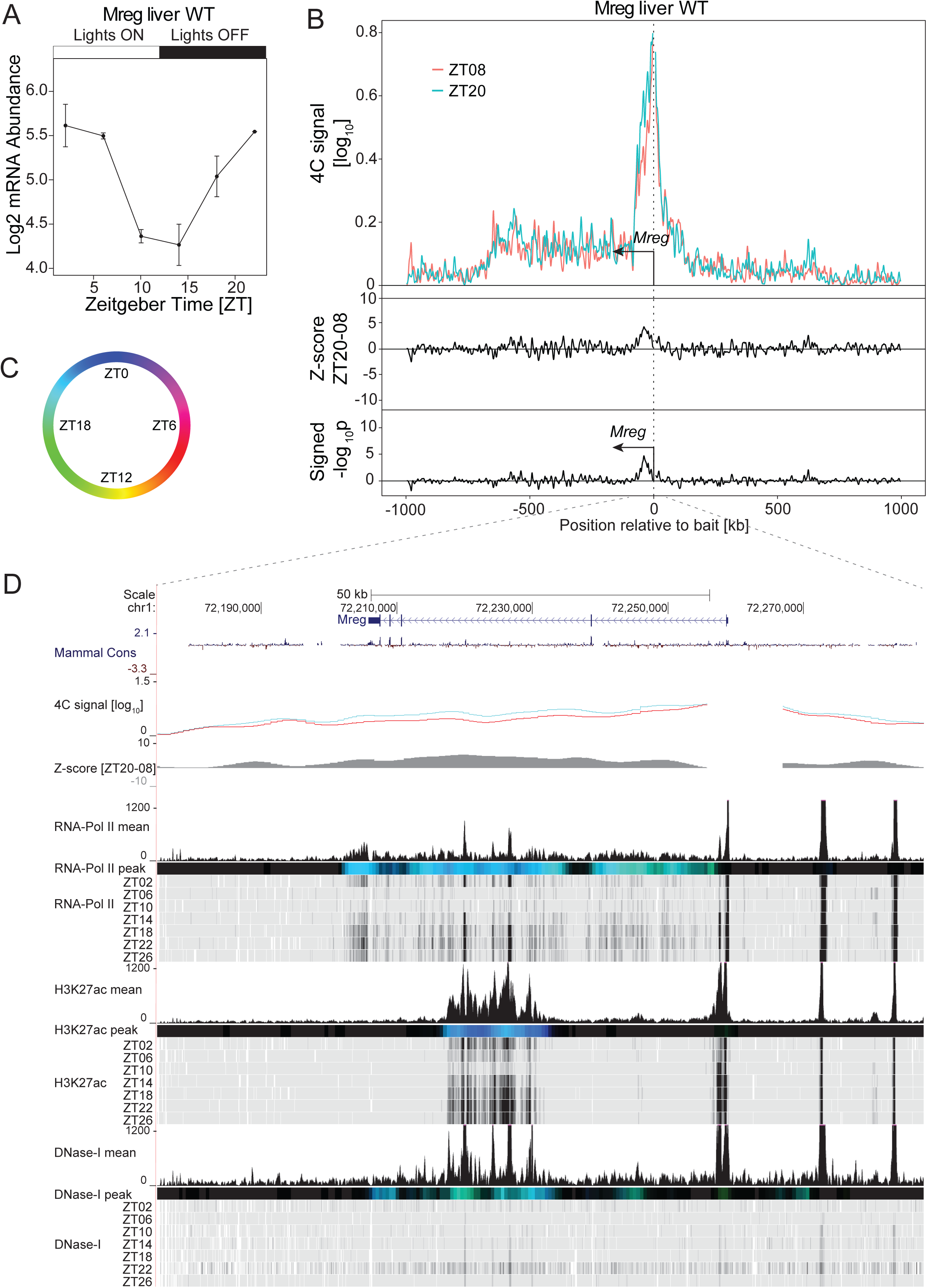
Oscillating interactions between the *Mreg* promoter and an intragenic enhancer-like elements. (A) *Mreg* mRNA expression over time in WT mouse liver (Atger et al.). (B) 4C-seq signal at ZT08 (red, n=2) and ZT20 (green, n=2) in WT mouse liver in a 2Mb genomic window surrounding the *Mreg* bait position (upper panel) and the corresponding Z-scores (middle track) and p-values (lower track) revealing a ZT20 preferential contact within a region located ∼40 kb downstream of the bait position. (C and D) Genome browser view with the *Mreg* 4C-seq signal at ZT08 (red) and ZT20 (blue) and time-resolved ChIP-seq signal for PolII, H3K27ac and DNase1 hypersensitivity in WT mouse liver (D) (Sobel et al.). Colored tracks represent peak time in chromatin marks following the color code as in (C) (methods). The interacting region located at ∼40 kb downstream of the *Mreg* TSS is marked by rhythmss in DHSs and H3K27ac peaking around ZT20, synchronously with *Mreg* transcription.

**Supplemental Figure 7.**
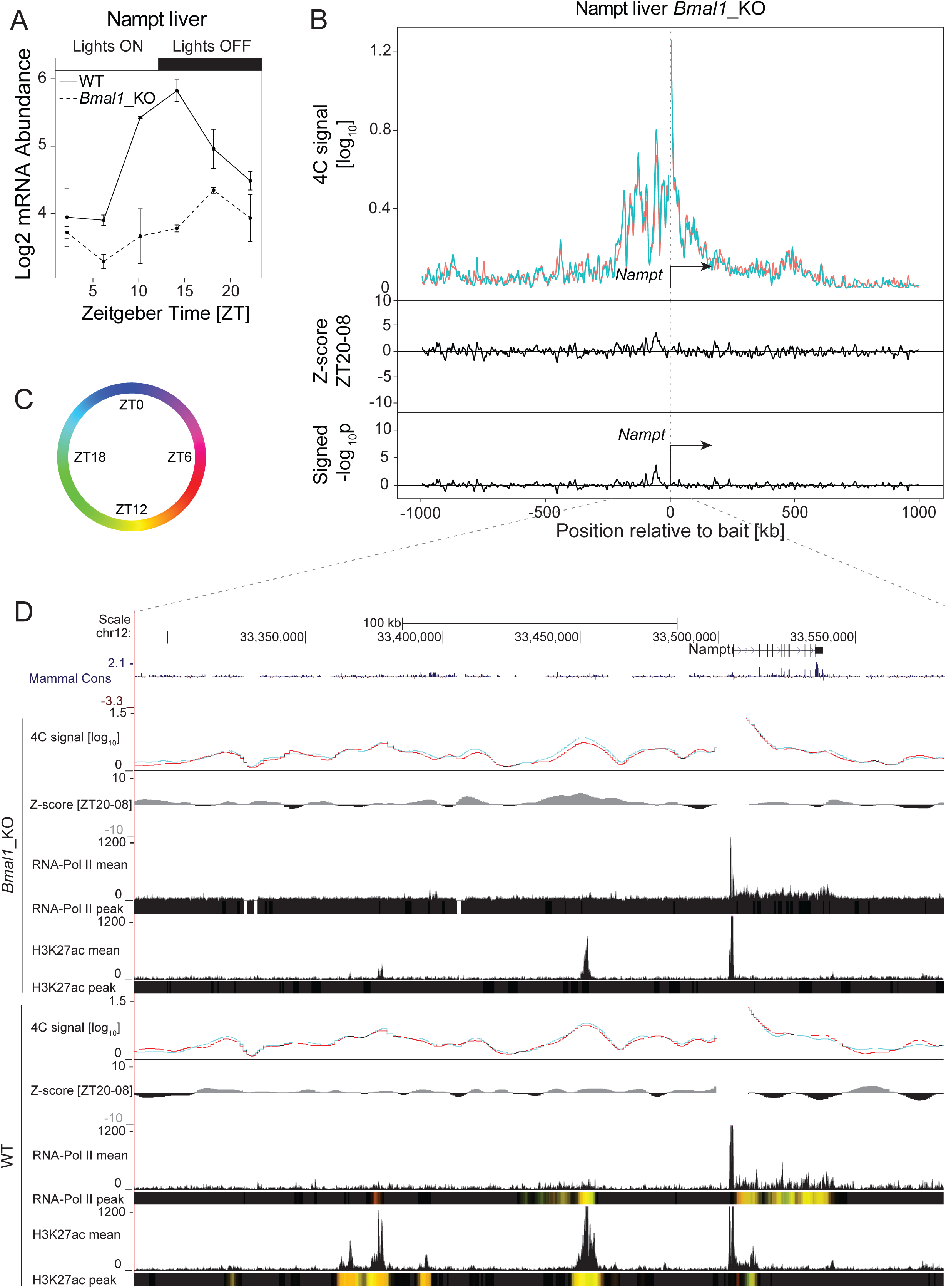
*Nampt* promoter-enhancer interactions do not depend on BMAL1. (A) *Nampt* transcript accumulation over time in WT (solid line) and *Bmal1* KO (dashed line) livers (Atger et al.). (B) *Nampt* 4C-seq signal at ZT08 (red, n=3) and ZT20 (green, n=3) in a genomic window of 2Mb surrounding the bait in the liver of *Bmal1* KO animals and the corresponding Z-scores and p-values. (C and D) Genome browser view of the *Nampt* 4C-seq signal and the corresponding Z-scores in livers of WT and *Bmal1* KO animals at ZT08 (red line) and ZT20 (blue line) (D). Mean and peak time of PolII and H3K27ac ChIP-seq signals are shown for both WT and *Bmal1* KO conditions (D) (Sobel et al.). Colored tracks represent peak time in chromatin marks following the color code as in (C) (methods). H3K27ac rhythms are observed at connected regions in WT livers. In the arrhythmic *Bmal1* KO livers, the chromatin marks no longer oscillate while chromatin contacts are maintained at levels comparable to WT conditions.

**Supplemental Figure 8.**
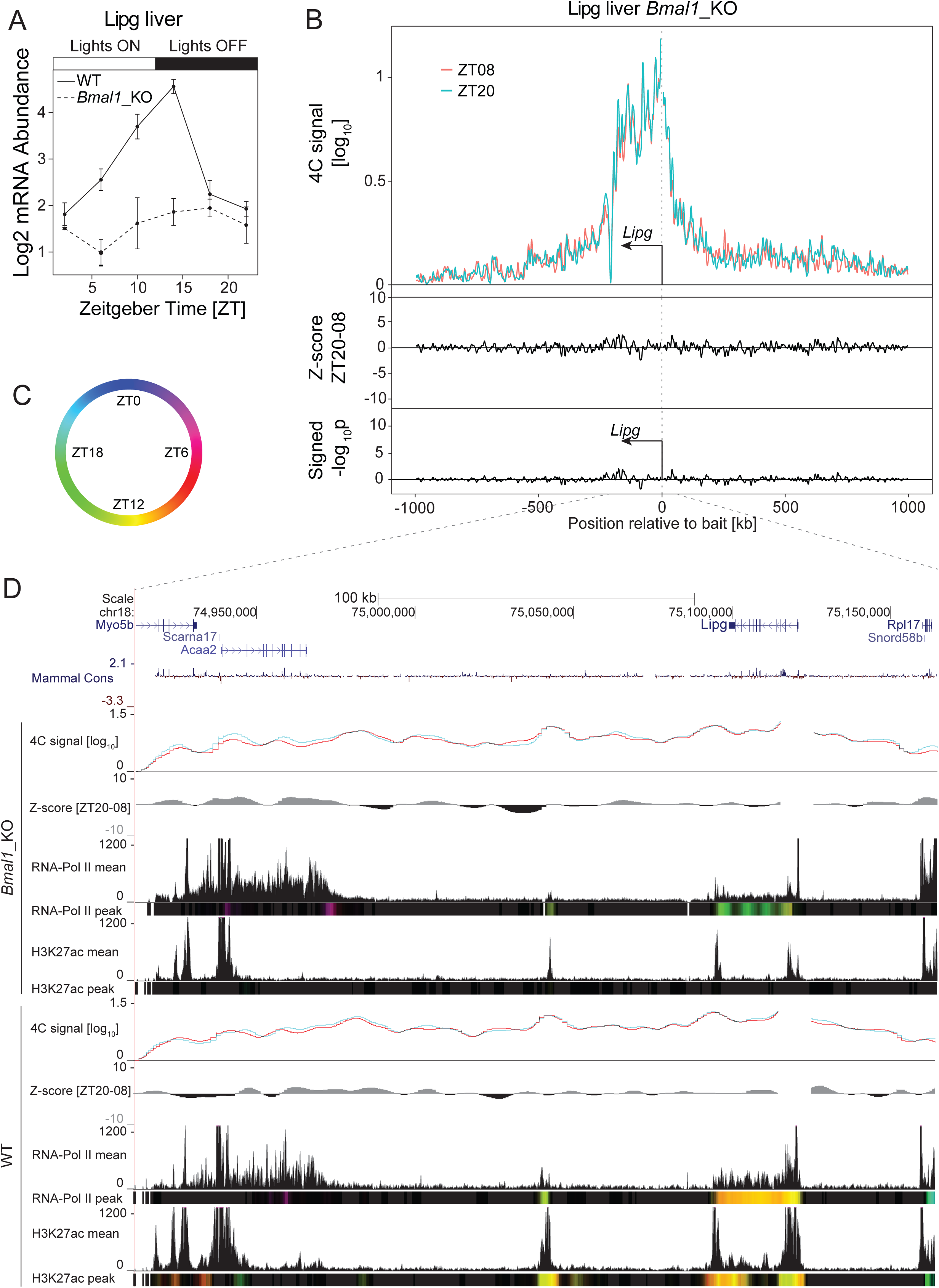
*Lipg* promoter-enhancer interactions do not depend on BMAL1. (A) *Lipg* transcripts accumulation over time in WT (black line) and *Bmal1* KO (grey line) livers (Atger et al.). (B) *Lipg* 4C-seq signal at ZT08 (red, n=3) and ZT20 (green, n=3) in a genomic window of 2Mb surrounding the bait in the liver of *Bmal1* KO animals and the corresponding Z-scores and p-values (C and D). Genome browser viewing of the *Lipg* 4C-seq signal and the corresponding Z-scores in livers of WT and *Bmal1* KO animals at ZT08 (red line) and ZT20 (blue line) (D). Mean and peak time of PolII and H3K27ac ChIP-seq signals are shown for both WT and *Bmal1* KO conditions (D) (Sobel et al.). Colored tracks represent peak time in chromatin marks following the color code as in (C) (methods). H3K27ac rhythms are observed at connected regions in WT livers. In the arrhythmic *Bmal1* KO livers, chromatin marks no longer oscillate while chromatin interactions are maintained at levels comparable to WT conditions.

**Supplementary Figure 9:**
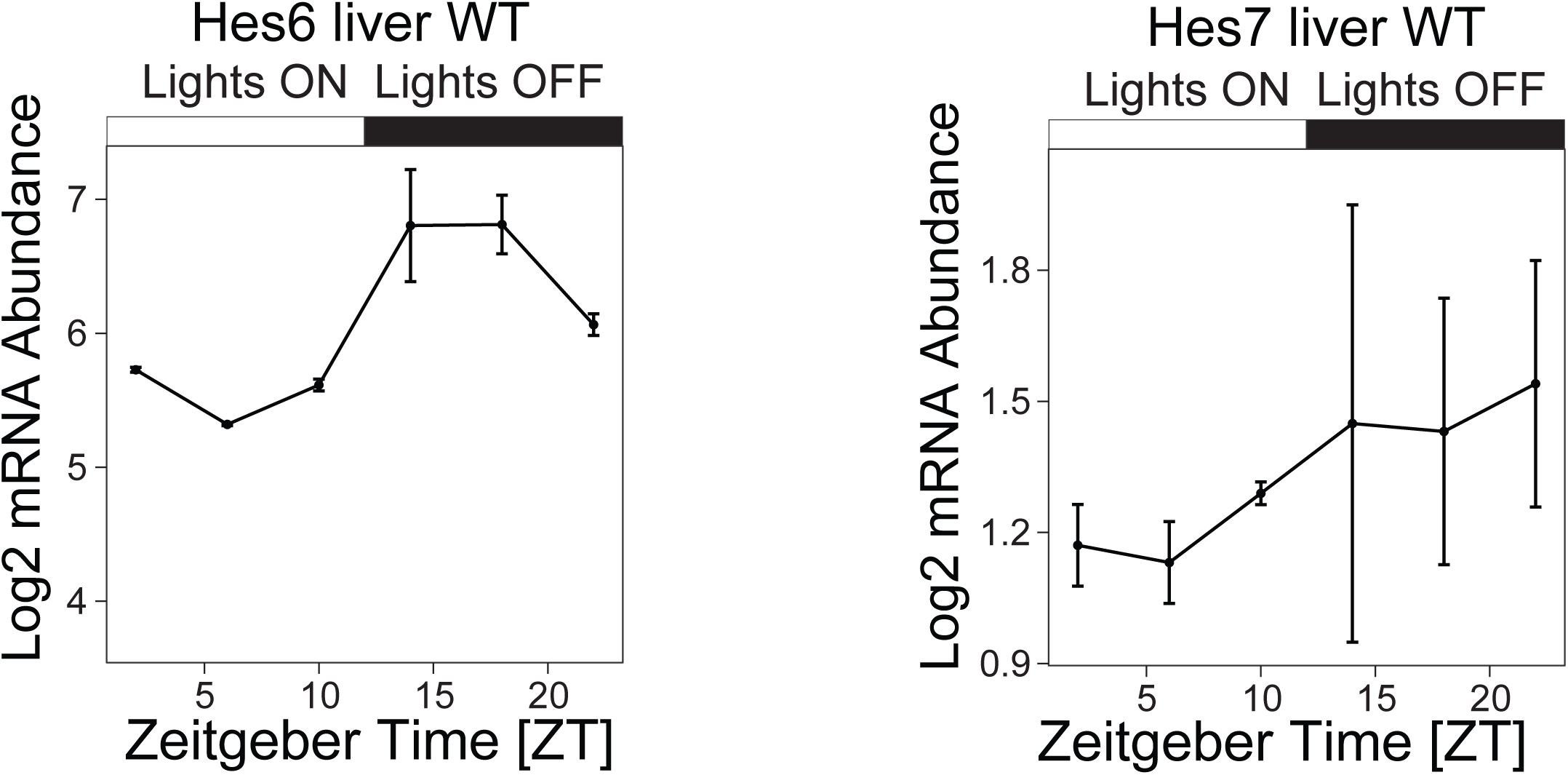
*Hes6* (left panel) and *Hes7* (right panel) transcript accumulation over time in WT livers. Data from (Atger et al.). *Hes6* mRNAs accumulate synchronously with *Per2* transcripts (Figure 2), and, despite low level of expression, the *Hes7* profile suggests a rhythmic expression in phase with *Per1* (Supplementary figure 1).

**Supplemental Figure 10.**
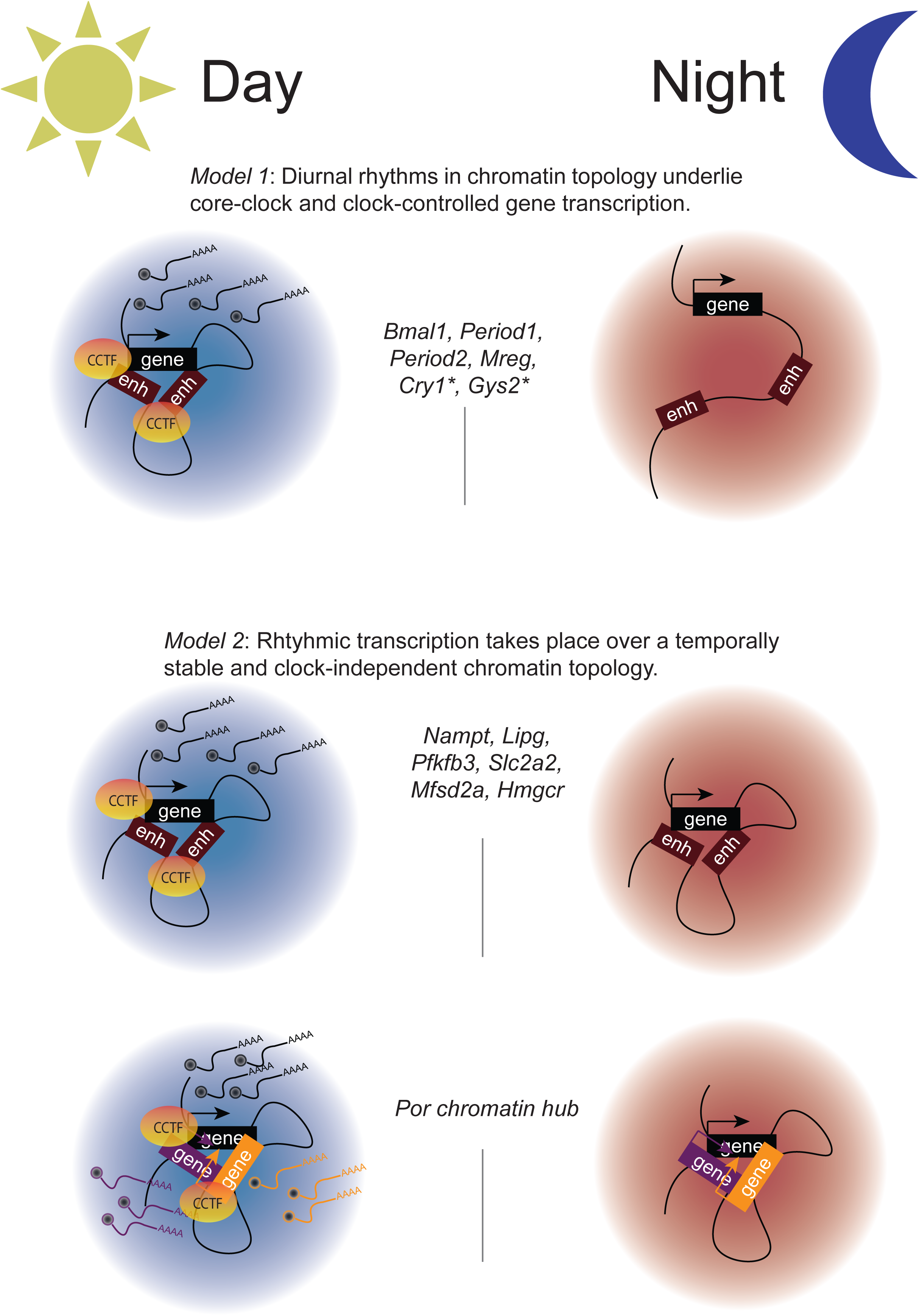
Models for diurnal transcription regulation in a 3-dimensional context of chromatin. *Model 1*: Dynamical interactions between oscillating gene promoter and surrounding regulatory elements accompany diurnal rhythms in transcription cycles. * = see (Mermet et al.) for a detailed analysis of these examples. *Model 2*: transcription oscillations take place over a static chromatin scaffold. CCTF: circadian clock transcription factors.

**Supplementary Table 1:** The table contains total 4C-sequencing read counts for each sample and the number and proportion of reads on *trans* and *cis* chromosome as well as on a 2Mb genomic region surrounding the bait fragment.

| Bait | Time | Tissue | Replicate | Passed Quality Threshold | Total Counts excluding +/- 5 fragments | Counts on Trans Chr | % of Total Counts on Trans Chr | Counts on Cis Chr | % of Total Counts on Cis Chr | Counts on Within 2Mb | % of Cis Counts Within 2Mb |
| --- | --- | --- | --- | --- | --- | --- | --- | --- | --- | --- | --- |
| HMGCR | ZT08 | WT | 1 | FALSE | 1035366 | 543345 | 52 | 492021 | 47 | 237617 | 48 |
| HMGCR | ZT08 | WT | 2 | FALSE | 3182892 | 1379151 | 43 | 1803740 | 56 | 941469 | 52 |
| HMGCR | ZT08 | WT | 3 | TRUE | 4771930 | 2207969 | 46 | 2563961 | 53 | 1194789 | 46 |
| HMGCR | ZT08 | WT | 4 | TRUE | 3716927 | 1965873 | 52 | 1751054 | 47 | 809875 | 46 |
| HMGCR | ZT20 | WT | 1 | TRUE | 3338601 | 1395036 | 41 | 1943564 | 58 | 1076131 | 55 |
| HMGCR | ZT20 | WT | 2 | TRUE | 3649876 | 1738302 | 47 | 1911574 | 52 | 918963 | 48 |
| HMGCR | ZT20 | WT | 3 | TRUE | 4449140 | 2069890 | 46 | 2379250 | 53 | 1219020 | 51 |
| HMGCR | ZT20 | WT | 4 | TRUE | 2661971 | 1147374 | 43 | 1514597 | 56 | 840986 | 55 |
| LIPG | ZT08 | WT | 1 | FALSE | 3563366 | 1513798 | 42 | 2049568 | 57 | 1263340 | 61 |
| LIPG | ZT08 | WT | 2 | TRUE | 2881818 | 1062206 | 36 | 1819612 | 63 | 1163180 | 63 |
| LIPG | ZT08 | WT | 3 | TRUE | 3913481 | 1553300 | 39 | 2360180 | 60 | 1459183 | 61 |
| LIPG | ZT08 | WT | 4 | TRUE | 3502098 | 1567320 | 44 | 1934778 | 55 | 1160060 | 59 |
| LIPG | ZT20 | WT | 1 | TRUE | 3340232 | 1151441 | 34 | 2188791 | 65 | 1478038 | 67 |
| LIPG | ZT20 | WT | 2 | TRUE | 4010230 | 1616122 | 40 | 2394107 | 59 | 1507873 | 62 |
| LIPG | ZT20 | WT | 3 | TRUE | 3333251 | 1221683 | 36 | 2111567 | 63 | 1372789 | 65 |
| LIPG | ZT20 | WT | 4 | TRUE | 4283620 | 1513163 | 35 | 2770456 | 64 | 1823188 | 65 |
| LIPG | ZT08 | KO | 1 | TRUE | 7024327 | 4974874 | 70 | 2049453 | 29 | 1223066 | 59 |
| LIPG | ZT08 | KO | 2 | TRUE | 4594070 | 2235352 | 48 | 2358718 | 51 | 1500426 | 63 |
| LIPG | ZT08 | KO | 3 | TRUE | 7513475 | 3014937 | 40 | 4498538 | 59 | 2912758 | 64 |
| LIPG | ZT20 | KO | 1 | TRUE | 7485984 | 5304137 | 70 | 2181847 | 29 | 1234124 | 56 |
| LIPG | ZT20 | KO | 2 | TRUE | 6602283 | 3270335 | 49 | 3331948 | 50 | 2232802 | 67 |
| LIPG | ZT20 | KO | 3 | TRUE | 7229646 | 2881566 | 39 | 4348080 | 60 | 2807011 | 64 |
| MFSD2A | ZT08 | WT | 1 | FALSE | 1717702 | 747381 | 43 | 970320 | 56 | 365875 | 37 |
| MFSD2A | ZT08 | WT | 2 | TRUE | 2930362 | 1001808 | 34 | 1928554 | 65 | 741595 | 38 |
| MFSD2A | ZT08 | WT | 3 | TRUE | 3133479 | 1136090 | 36 | 1997389 | 63 | 754194 | 37 |
| MFSD2A | ZT08 | WT | 4 | TRUE | 2585981 | 1140890 | 44 | 1445091 | 55 | 525149 | 36 |
| MFSD2A | ZT20 | WT | 1 | TRUE | 3660025 | 1167515 | 31 | 2492509 | 68 | 1048781 | 42 |
| MFSD2A | ZT20 | WT | 2 | TRUE | 3279541 | 1230645 | 37 | 2048896 | 62 | 771063 | 37 |
| MFSD2A | ZT20 | WT | 3 | TRUE | 2518377 | 940536 | 37 | 1577841 | 62 | 654171 | 41 |
| MFSD2A | ZT20 | WT | 4 | FALSE | 2196529 | 672723 | 30 | 1523806 | 69 | 641237 | 42 |
| Mreg | ZT08 | Liver | 1 | TRUE | 1450572 | 509636 | 35 | 940936 | 64 | 382708 | 40 |
| Mreg | ZT08 | Liver | 2 | TRUE | 2255183 | 844506 | 37 | 1410676 | 62 | 551107 | 39 |
| Mreg | ZT20 | Liver | 1 | TRUE | 1498447 | 502640 | 33 | 995807 | 66 | 448571 | 45 |
| Mreg | ZT20 | Liver | 2 | TRUE | 1699239 | 574620 | 33 | 1124619 | 66 | 518165 | 46 |
| NAMPT | ZT08 | WT | 1 | FALSE | 1583818 | 818302 | 51 | 765516 | 48 | 378293 | 49 |
| NAMPT | ZT08 | WT | 2 | FALSE | 1948085 | 801050 | 41 | 1147035 | 58 | 600793 | 52 |
| NAMPT | ZT08 | WT | 3 | TRUE | 3015387 | 1280207 | 42 | 1735179 | 57 | 881520 | 50 |
| NAMPT | ZT08 | WT | 4 | TRUE | 1925480 | 971734 | 50 | 953746 | 49 | 470348 | 49 |
| NAMPT | ZT20 | WT | 1 | TRUE | 2503653 | 920727 | 36 | 1582926 | 63 | 933947 | 59 |
| NAMPT | ZT20 | WT | 2 | TRUE | 3355484 | 1449046 | 43 | 1906438 | 56 | 997131 | 52 |
| NAMPT | ZT20 | WT | 3 | TRUE | 2327322 | 903755 | 38 | 1423567 | 61 | 797718 | 56 |
| NAMPT | ZT20 | WT | 4 | TRUE | 1181151 | 392118 | 33 | 789032 | 66 | 461475 | 58 |
| NAMPT | ZT08 | KO | 1 | TRUE | 3503168 | 2776746 | 79 | 726421 | 20 | 331090 | 45 |
| NAMPT | ZT08 | KO | 2 | TRUE | 1392323 | 834781 | 59 | 557542 | 40 | 284874 | 51 |
| NAMPT | ZT08 | KO | 3 | TRUE | 4535535 | 2270260 | 50 | 2265275 | 49 | 1186780 | 52 |
| NAMPT | ZT20 | KO | 1 | TRUE | 3972957 | 3223949 | 81 | 749008 | 18 | 329104 | 43 |
| NAMPT | ZT20 | KO | 2 | TRUE | 3586455 | 2137346 | 59 | 1449109 | 40 | 813814 | 56 |
| NAMPT | ZT20 | KO | 3 | TRUE | 3608858 | 1828525 | 50 | 1780333 | 49 | 942469 | 52 |
| PFKFB3 | ZT08 | WT | 1 | FALSE | 5527841 | 2401730 | 43 | 3126111 | 56 | 1569812 | 50 |
| PFKFB3 | ZT08 | WT | 2 | TRUE | 5902530 | 1824760 | 30 | 4077770 | 69 | 2238633 | 54 |
| PFKFB3 | ZT08 | WT | 3 | TRUE | 4704636 | 1603483 | 34 | 3101153 | 65 | 1650888 | 53 |
| PFKFB3 | ZT08 | WT | 4 | TRUE | 4050685 | 1751602 | 43 | 2299083 | 56 | 1156191 | 50 |
| PFKFB3 | ZT20 | WT | 1 | TRUE | 3576186 | 1026471 | 28 | 2549715 | 71 | 1515692 | 59 |
| PFKFB3 | ZT20 | WT | 2 | TRUE | 4084536 | 1411092 | 34 | 2673444 | 65 | 1468369 | 54 |
| PFKFB3 | ZT20 | WT | 3 | TRUE | 5946705 | 1962108 | 32 | 3984597 | 67 | 2323935 | 58 |
| PFKFB3 | ZT20 | WT | 4 | TRUE | 5779292 | 1495807 | 25 | 4283484 | 74 | 2565147 | 59 |
| POR | ZT08 | WT | 1 | FALSE | 5111888 | 2452457 | 47 | 2659430 | 52 | 1487092 | 55 |
| POR | ZT08 | WT | 2 | TRUE | 5533160 | 2326356 | 42 | 3206804 | 57 | 1889951 | 58 |
| POR | ZT08 | WT | 3 | TRUE | 4687742 | 2187205 | 46 | 2500537 | 53 | 1403425 | 56 |
| POR | ZT08 | WT | 4 | TRUE | 3687557 | 1953166 | 52 | 1734390 | 47 | 967233 | 55 |
| POR | ZT20 | WT | 1 | TRUE | 5050192 | 2016007 | 39 | 3034185 | 60 | 1874307 | 61 |
| POR | ZT20 | WT | 2 | TRUE | 4102235 | 1946566 | 47 | 2155669 | 52 | 1227763 | 56 |
| POR | ZT20 | WT | 3 | TRUE | 5166265 | 2323553 | 44 | 2842711 | 55 | 1708327 | 60 |
| POR | ZT20 | WT | 4 | TRUE | 5304647 | 2066206 | 38 | 3238441 | 61 | 2001630 | 61 |
| POR | ZT08 | KO | 1 | TRUE | 6582610 | 3950107 | 60 | 2632502 | 39 | 1530391 | 58 |
| POR | ZT08 | KO | 2 | TRUE | 4349106 | 2598421 | 59 | 1750685 | 40 | 967491 | 55 |
| POR | ZT08 | KO | 3 | TRUE | 4317498 | 2210032 | 51 | 2107465 | 48 | 1239674 | 58 |
| POR | ZT20 | KO | 1 | TRUE | 4612843 | 3242029 | 70 | 1370814 | 29 | 735100 | 53 |
| POR | ZT20 | KO | 2 | TRUE | 4573065 | 2739142 | 59 | 1833923 | 40 | 1122115 | 61 |
| POR | ZT20 | KO | 3 | TRUE | 2868922 | 1461655 | 50 | 1407267 | 49 | 825727 | 58 |
| SLC2A2 | ZT08 | WT | 1 | FALSE | 3577101 | 1637980 | 45 | 1939121 | 54 | 835495 | 43 |
| SLC2A2 | ZT08 | WT | 2 | FALSE | 3798188 | 1356961 | 35 | 2441227 | 64 | 1089889 | 44 |
| SLC2A2 | ZT08 | WT | 3 | TRUE | 4019514 | 1527120 | 37 | 2492394 | 62 | 1135285 | 45 |
| SLC2A2 | ZT08 | WT | 4 | TRUE | 3094019 | 1424636 | 46 | 1669382 | 53 | 766702 | 45 |
| SLC2A2 | ZT20 | WT | 1 | TRUE | 4645041 | 1559712 | 33 | 3085328 | 66 | 1575793 | 51 |
| SLC2A2 | ZT20 | WT | 2 | TRUE | 3579413 | 1354758 | 37 | 2224654 | 62 | 1069270 | 48 |
| SLC2A2 | ZT20 | WT | 3 | TRUE | 3534188 | 1267346 | 35 | 2266842 | 64 | 1101578 | 48 |
| SLC2A2 | ZT20 | WT | 4 | TRUE | 2839046 | 926911 | 32 | 1912135 | 67 | 988630 | 51 |
| Bmal1_TSS | ZT00 | WT | 1 | TRUE | 1079899 | 370562 | 34 | 709337 | 65 | 354302 | 49 |
| Bmal1_TSS | ZT00 | WT | 2 | TRUE | 1409403 | 510271 | 36 | 899132 | 63 | 428552 | 47 |
| Bmal1_TSS | ZT04 | WT | 1 | TRUE | 1744960 | 534439 | 30 | 1210521 | 69 | 636046 | 52 |
| Bmal1_TSS | ZT04 | WT | 2 | FALSE | 1428276 | 457631 | 32 | 970644 | 67 | 506502 | 52 |
| Bmal1_TSS | ZT08 | WT | 1 | TRUE | 1511241 | 495839 | 32 | 1015401 | 67 | 533286 | 52 |
| Bmal1_TSS | ZT08 | WT | 2 | TRUE | 1483911 | 537711 | 36 | 946200 | 63 | 494382 | 52 |
| Bmal1_TSS | ZT12 | WT | 1 | TRUE | 1529470 | 471147 | 30 | 1058323 | 69 | 502279 | 47 |
| Bmal1_TSS | ZT12 | WT | 2 | FALSE | 1638928 | 622879 | 38 | 1016049 | 61 | 465479 | 45 |
| Bmal1_TSS | ZT16 | WT | 1 | TRUE | 1401632 | 437628 | 31 | 964003 | 68 | 511289 | 53 |
| Bmal1_TSS | ZT16 | WT | 2 | TRUE | 1343072 | 442328 | 32 | 900744 | 67 | 466333 | 51 |
| Bmal1_TSS | ZT20 | WT | 1 | TRUE | 1285656 | 491020 | 38 | 794636 | 61 | 429430 | 54 |
| Bmal1_TSS | ZT20 | WT | 2 | FALSE | 1430674 | 532027 | 37 | 898647 | 62 | 413146 | 45 |
| Per1_TSS | ZT00 | WT | 1 | TRUE | 1175883 | 499210 | 42 | 676673 | 57 | 291024 | 43 |
| Per1_TSS | ZT00 | WT | 2 | TRUE | 1000482 | 437007 | 43 | 563475 | 56 | 245944 | 43 |
| Per1_TSS | ZT04 | WT | 1 | TRUE | 1000658 | 387195 | 38 | 613463 | 61 | 254586 | 41 |
| Per1_TSS | ZT04 | WT | 2 | TRUE | 1135041 | 487754 | 42 | 647287 | 57 | 259955 | 40 |
| Per1_TSS | ZT08 | WT | 1 | TRUE | 1049270 | 432660 | 41 | 616610 | 58 | 284276 | 46 |
| Per1_TSS | ZT08 | WT | 2 | TRUE | 1138240 | 518759 | 45 | 619481 | 54 | 298999 | 48 |
| Per1_TSS | ZT12 | WT | 1 | TRUE | 917112 | 377550 | 41 | 539561 | 58 | 250308 | 46 |
| Per1_TSS | ZT12 | WT | 2 | TRUE | 1003821 | 435892 | 43 | 567929 | 56 | 242225 | 42 |
| Per1_TSS | ZT16 | WT | 1 | TRUE | 908334 | 345520 | 38 | 562813 | 61 | 264559 | 47 |
| Per1_TSS | ZT16 | WT | 2 | TRUE | 1110747 | 451209 | 40 | 659538 | 59 | 335630 | 50 |
| Per1_TSS | ZT20 | WT | 1 | TRUE | 1000583 | 500730 | 50 | 499852 | 49 | 194284 | 38 |
| Per1_TSS | ZT20 | WT | 2 | TRUE | 1063557 | 432035 | 40 | 631522 | 59 | 288402 | 45 |
| Per2_TSS | ZT00 | WT | 1 | TRUE | 1067769 | 358552 | 33 | 709216 | 66 | 416503 | 58 |
| Per2_TSS | ZT00 | WT | 2 | TRUE | 1227622 | 426132 | 34 | 801490 | 65 | 463091 | 57 |
| Per2_TSS | ZT04 | WT | 1 | TRUE | 1646936 | 543315 | 32 | 1103621 | 67 | 595390 | 53 |
| Per2_TSS | ZT04 | WT | 2 | TRUE | 1184481 | 432994 | 36 | 751487 | 63 | 399351 | 53 |
| Per2_TSS | ZT08 | WT | 1 | TRUE | 1164352 | 368606 | 31 | 795746 | 68 | 440422 | 55 |
| Per2_TSS | ZT08 | WT | 2 | TRUE | 1156916 | 440267 | 38 | 716648 | 61 | 402921 | 56 |
| Per2_TSS | ZT12 | WT | 1 | TRUE | 1577594 | 480105 | 30 | 1097489 | 69 | 583207 | 53 |
| Per2_TSS | ZT12 | WT | 2 | TRUE | 1151334 | 413899 | 35 | 737434 | 64 | 368952 | 50 |
| Per2_TSS | ZT16 | WT | 1 | TRUE | 1453417 | 453258 | 31 | 1000158 | 68 | 610203 | 61 |
| Per2_TSS | ZT16 | WT | 2 | TRUE | 1065073 | 358388 | 33 | 706685 | 66 | 441241 | 62 |
| Per2_TSS | ZT20 | WT | 1 | TRUE | 1395584 | 508545 | 36 | 887039 | 63 | 452851 | 51 |
| Per2_TSS | ZT20 | WT | 2 | TRUE | 1316675 | 456211 | 34 | 860464 | 65 | 474896 | 55 |

**Supplementary Table 2:**
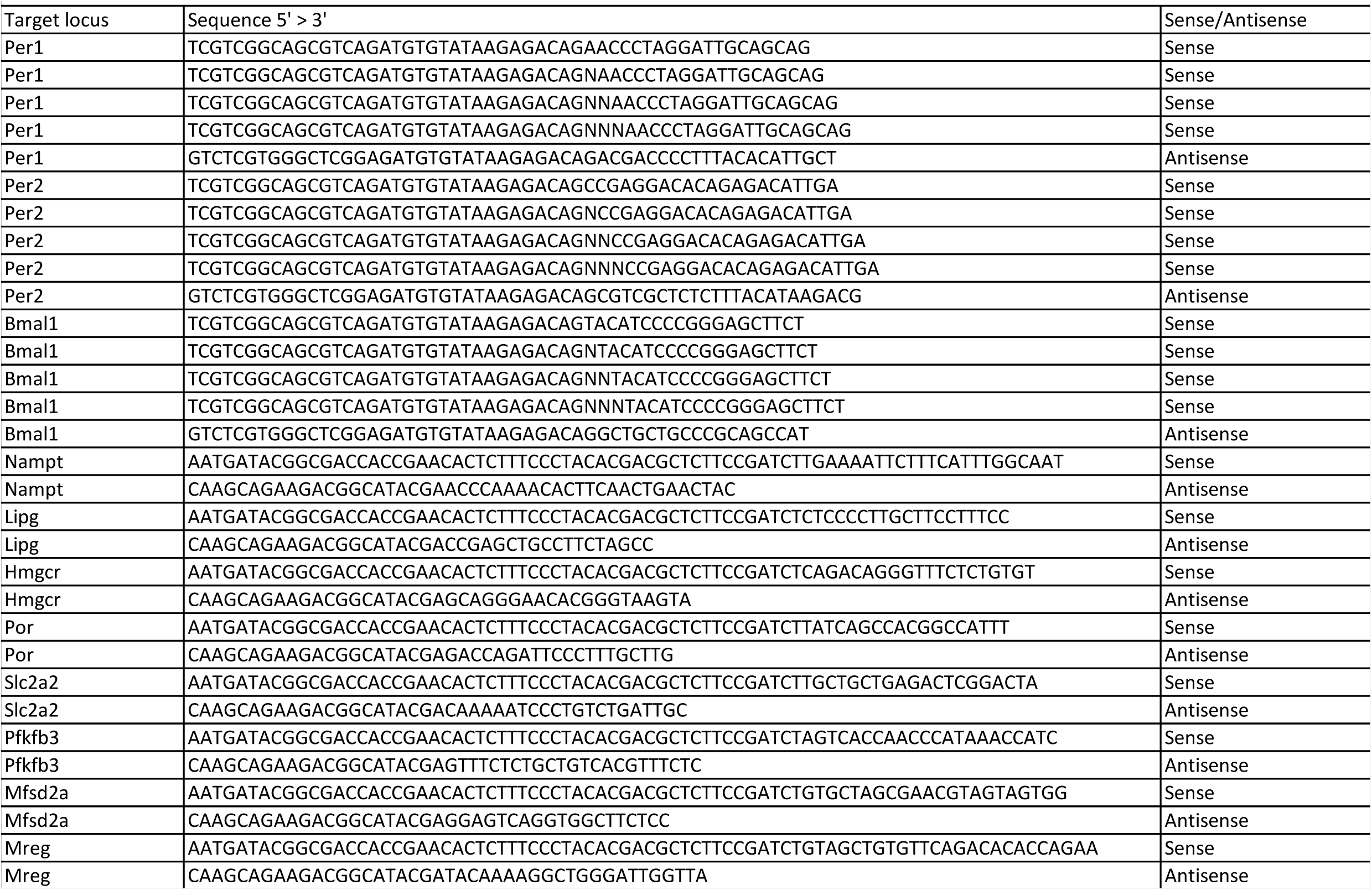
Table contains sequence of inverse-PCR primers used to generate 4C-seq libraries.

